# Chemical stimuli override a temperature-dependent morphological program by reprogramming the transcriptome of a fungal pathogen

**DOI:** 10.1101/2023.04.21.537729

**Authors:** Dror Assa, Mark Voorhies, Anita Sil

**Affiliations:** Department of Microbiology and Immunology, University of California San Francisco, San Francisco, California, United States of America

## Abstract

The human fungal pathogen *Histoplasma* changes its morphology in response to temperature. At 37°C it grows as a budding yeast whereas at room temperature it transitions to hyphal growth. Prior work has demonstrated that 15-20% of transcripts are temperature-regulated, and that transcription factors Ryp1-4 are necessary to establish yeast growth. However, little is known about transcriptional regulators of the hyphal program. To identify TFs that regulate filamentation, we utilize chemical inducers of hyphal growth. We show that addition of cAMP analogs or an inhibitor of cAMP breakdown overrides yeast morphology, yielding inappropriate hyphal growth at 37°C. Additionally, butyrate supplementation triggers hyphal growth at 37°C. Transcriptional profiling of cultures filamenting in response to cAMP or butyrate reveals that a limited set of genes respond to cAMP while butyrate dysregulates a larger set. Comparison of these profiles to previous temperature- or morphology-regulated gene sets identifies a small set of morphology-specific transcripts. This set contains 9 TFs of which we characterized three, *STU1*, *FBC1*, and *PAC2*, whose orthologs regulate development in other fungi. We found that each of these TFs is individually dispensable for room-temperature (RT) induced filamentation but each is required for other aspects of RT development. *FBC1* and *PAC2*, but not *STU1*, are necessary for filamentation in response to cAMP at 37°C. Ectopic expression of each of these TFs is sufficient to induce filamentation at 37°C. Finally, *PAC2* induction of filamentation at 37°C is dependent on *STU1*, suggesting these TFs form a regulatory circuit that, when activated at RT, promotes the hyphal program.

**Importance:** Fungal illnesses pose a significant disease burden. However, the regulatory circuits that govern the development and virulence of fungi remain largely unknown. This study utilizes chemicals that can override the normal growth morphology of the human pathogen *Histoplasma*. Using transcriptomic approaches, we identify novel regulators of hyphal morphology and refine our understanding of the transcriptional circuits governing morphology in *Histoplasma*.

## Introduction

To survive in diverse environments, microbes employ a variety of mechanisms to sense and respond to distinct signals, allowing them to adapt to different conditions. Thermally dimorphic fungi have developed the ability to thrive in diverse conditions, growing as hyphae at ambient temperatures and at room temperature (RT) in the laboratory, but switching to a unicellular growth form at mammalian body temperature (37°C in the laboratory). *Histoplasma capsulatum* is one such thermally dimorphic fungus that grows as sporulating hyphae in the soil and, after inhalation, changes its shape and gene expression program to generate virulent yeast cells in response to mammalian body temperature. This transition to yeast cells can result in an acute pulmonary infection known as histoplasmosis (1, 2). The process of morphological differentiation requires the ability to integrate signals on a molecular level and convert those signals into transcriptional, translational and biochemical changes that orchestrate the transition from one growth form to another.

The molecular basis of *Histoplasma’s* ability to transition between morphologies has been at the focus of many studies in the past decades. Previous work had shown that 15-20% of transcripts in *Histoplasma* are regulated by temperature (3–5). Specific functions, such as virulence, iron acquisition, and cell wall modification are correlated with 37°C and yeast-phase growth, whereas the hyphal growth form that dominates at room temperature promotes the expression of enzymes such as tyrosinases, cytochrome p450s, oxidoreducatases, and peroxidases. Additionally, the abundance of *Histoplasma* transcripts encoding 18 putative transcription factors is increased in hyphae compared to yeast (3).

Forward genetic screens have been instrumental in finding regulators of morphology, identifying Drk1, a hybrid histidine kinase orthologous to *Candida albicans* Nik1 (6), which is necessary for yeast morphology (7), and 3 transcription factors that are also required for yeast phase (RYP) growth: Ryp1, a WOPR family transcription factor orthologous to *Candida albicans* Wor1, and Ryp2/3, both velvet family transcription factors orthologous to *Aspergillus* VosA and VelB, respectively (8, 9). Follow up studies also identified an additional transcription factor required for yeast phase growth, Ryp4, a Zn(II)2Cys6 zinc binuclear cluster domain protein (4). Furthermore, Gilmore *et al.* showed that inappropriate expression of the transcription factor Wet1 is sufficient to cause filamentous growth at 37°C in *Histoplasma* (3). Wet1 is orthologous to wetA in *Aspergillus*, which regulates the late stages of asexual spore production (10). More recently, Rodriguez *et al.* reported that expression of the APSES-family transcription factor Stu1, which is orthologous to StuA in *Aspergillus* and Enhanced Filamentous Growth 1 (Efg1) in *C. albicans*, is also sufficient to trigger inappropriate filamentous growth at 37°C in *Histoplasma* (11). Additionally, Longo et al showed that using RNA interference to target Stu1 resulted in decreased formation of aerial hyphae on plates (12). StuA coordinates conidiophore formation in the *Aspergilli* (13, 14), and Efg1 is necessary for filamentous growth in *C. albicans* (reviewed in (15)).

Here we examine the relationship between transcription, *Histoplasma* hyphal formation, and the cAMP/protein kinase A (PKA) pathway, which is a central and conserved signaling pathway in all eukaryotes. In fungi, the cAMP-PKA pathway regulates carbon utilization, mating, morphology, stress response, and virulence. The cAMP-PKA pathway has been the subject of extensive research in *S. cerevisiae* (reviewed in (16–18)). In this organism, fermentable carbon sources such as glucose are sensed by several pathways that lead to the activation of adenylate cyclase, which synthesizes cAMP from ATP. cAMP then binds the regulatory subunits of protein kinase A (PKA), releasing the catalytically active subunits to phosphorylate target proteins. One of the main roles of the cAMP-PKA pathway in *S. cerevisiae* is to allow rapid response to stress. In non-stress conditions, the transcription factors Msn2/4p are phosphorylated by PKA, preventing them from entering the cell nucleus (19). Various stress signals inactivate PKA rapidly and cause the induction of the environmental stress response (20–24). Other outcomes of PKA inactivation include induction of gluconeogenesis, respiration, and alternative energy source utilization pathways and repression of cell division and ribosomal biogenesis pathways (25, 26). The cAMP-PKA pathway also regulates pseudohyphal growth by activating the transcription factor Flo8p (27, 28). In *Candida* albicans, the cAMP-PKA pathway integrates signals such as environmental pH, N-acetylglucosamine (GlcNAc), and the presence of serum in growth media, and, depending on specific contexts, positively regulates white-opaque switching or the transition to hyphal growth, mediated by the activation of Efg1 (29–36). The activity of the cAMP-PKA pathway is also necessary for the expression of virulence genes in *C. albicans* (34, 37, 38). In *Aspergillus* spp., sensing of signals such as glucose by hetero-trimeric G-protein leads to activation of the cAMP-PKA pathway, inducing spore germination, inhibiting sporulation and coordinating the synthesis of secondary metabolites (39–44). Finally, intracellular and extracellular cAMP levels rise when *Histoplasma* G217B yeast (growing at 37°C) are shifted to room temperature, concurrent with the formation of hyphae (45). These data suggest a correlation between cAMP accumulation and filamentation. Moreover, addition of the stable cAMP analog dibutyryl-cAMP (dbcAMP) to *Histoplasma* cultures growing at 37°C promotes filamentous growth (45–47). Similarly, in the closely related dimorphic fungi *Paracoccidioides brasiliensis* and *Blastomyces dermatitidis*, exogenous cAMP delays the transition from hyphae to yeast (45, 46). However, the underlying molecular mechanisms of filamentation induced by exogenous activation of the cAMP pathway in thermally dimorphic fungi have not been examined.

In this study, we used transcriptional profiling of Histoplasma to gain insight to the molecular events that occur upon chemical induction of filamentation. By comparing these expression profiling experiments to RT-induced expression, we identified groups of transcripts whose accumulation is associated with yeast or hyphal morphologies. Finally, we focused on 3 transcription factors that have been shown to regulate morphology and development in other organisms and showed that they are capable of driving a hyphal program in *Histoplasma*.

## Results

### cAMP analogs drive inappropriate filamentation at 37°C

To characterize the effect of dbcAMP on *Histoplasma* growth, we added dbcAMP to early-log phase G217B and G217B*ura5* (WU15) cultures growing at 37°C in HMM containing glucose or N-acetylglucosamine (HMM+GlcNAc) as main carbon sources (Fig. 1A). Our previous work showed that GlcNAc stimulates hyphal growth upon shift from 37°C to RT (48). While no morphological difference was observed after one day, within 2 days a noticeable increase in filamentous growth occurred in cultures growing in HMM+GlcNAc and treated with dbcAMP compared to the vehicle (water). In contrast, 10 mM dbcAMP did not affect the morphology of yeast cultures in HMM containing only glucose (Fig. 1B). Hyphal growth was even more evident by the third day after dbcAMP addition (Fig. S1A). The extent to which dbcAMP promotes filamentation was concentration-dependent: a visible increase in filamentous growth could be observed in cultures with as little as 1.25 mM dbcAMP (Fig. S1B). Addition of dbcAMP to solid media (HMM agarose containing GlcNAc) also strongly favored filamentous growth at 37°C (Fig. 1C).

**Figure 1.**
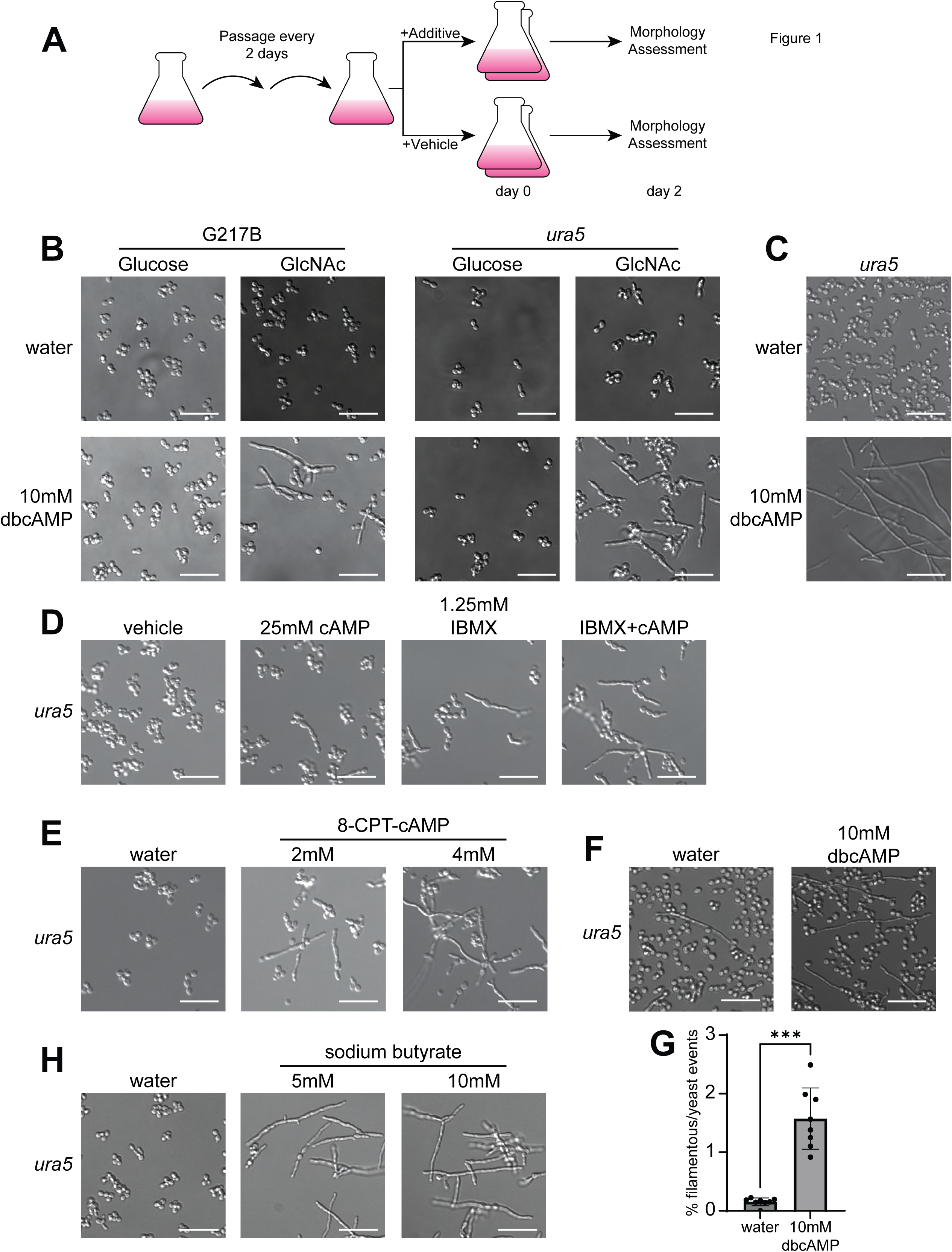
Exogenous cAMP promotes filamentous growth at 37°C. (A) Schematic outline of experiments testing the response of *Histoplasma* to chemical additives. (B) Cell morphology of G217B and G217B*ura5 Histoplasma* after 2 days of growth at 37°C in liquid HMM containing glucose or GlcNAc with or without 10mM dbcAMP. (C) Cell morphology of G217B*ura5 Histoplasma* after 8 days of growth on HMM-agarose plates containing GlcNAc, with or without 10mM dbcAMP, at 37°C. (D) Cell morphology of G217B*ura5 Histoplasma* after 2 days of growth at 37°C in liquid HMM containing GlcNAc with 1.25mM IBMX, 25mM cAMP, IBMX+cAMP, or water as vehicle control. (E) Cell morphology of G217B*ura5 Histoplasma* after 2 days of growth at 37°C in liquid HMM containing GlcNAc with 4mM or 2mM 8-CPT-cAMP, or water as vehicle control. (F) Cell morphology of G217B*ura5 Histoplasma* after 7 days of growth at RT in liquid HMM containing glucose with or without 10mM dbcAMP. (G) Quantification of filamentation after 7 days at RT in liquid HMM containing glucose with or without 10mM dbcAMP; N > 243 events. (H) Cell morphology of G217B*ura5 Histoplasma* after 2 days of growth at 37°C in liquid HMM containing GlcNAc with 5mM or 10mM sodium butyrate, or water as vehicle. Scale bar denotes 10 µm; *** p < 0.001, Wilcoxon rank-sum test.

We next wanted to test whether other means of increasing cAMP levels had a similar effect on hyphal formation. Normally cAMP is broken down within the cell by the action of phosphodiesterases. We attempted to raise cAMP levels by adding cAMP sodium salt or the phosphodiesterase inhibitor IBMX. Whereas 25 mM cAMP had a negligible effect on morphology on its own, the addition of 1.25 mM IBMX was sufficient to trigger filamentation within 2 days (Fig. 1D). Adding both cAMP and IBMX had a stronger filamentation effect, suggesting that minimizing the breakdown of exogenous cAMP by inhibiting phosphodiesterase activity may cause a net increase in cAMP levels that promotes filamentation. Furthermore, adding 8-(4-chlorophenylthio)adenosine 3’,5’-cyclic monophosphate (8-CPT-cAMP), an alternative cAMP analog, also promoted robust filamentation (Fig. 1E). Taken together, these data confirm that exogenously added cAMP overrides yeast phase growth and promotes filamentation at 37°C.

The transition from yeast to hyphae in response to temperature shift is less robust in glucose relative to GlcNAc (48). To determine if the addition of exogenous cAMP addition promotes filamentation in response to temperature shift in glucose, where the transition is asynchronous and inefficient, we added 10 mM dbcAMP to cultures shifted from 37°C to RT in glucose-containing media. We observed a subtle (Fig. 1F) but statistically significant (Fig. 1G) increase in elongated cells and filamentous structures in liquid culture by day 7 after shift in the presence of dbcAMP.

Since dbcAMP hydrolysis can release butyrate into the growth media, we also investigated the effect of adding sodium butyrate on its own to *Histoplasma* growth media. In the presence of GlcNAc, butyrate strongly induced filamentation within 2 days; however, the effect of butyrate was much more potent than dbcAMP. For example, 10 mM dbcAMP stimulated robust filamentation by day 2 (Fig. 1B), but yeast cells were still present. In contrast, 5 mM butyrate stimulated complete hyphal transformation with no yeast cells remaining in the cultures (Fig. 1H).

### cAMP analogs and butyrate stimulate robust transcriptional changes

To further understand the molecules and pathways that mediate filamentation after addition of cAMP analogs, we performed transcriptional profiling of G217B*ura5 Histoplasma* cultures grown in HMM+GlcNAc media at 37°C with addition of 10 mM dbcAMP, 4 mM 8-CPT-cAMP, 5mM butyrate, or water as vehicle control. Each of these additions stimulated filamentation compared to the water control (Fig. 2A). We found that 721 or 1083 transcripts were significantly up-regulated in response to dbcAMP or 8-CPT-cAMP, respectively (>1.5 fold change, FDR < 5%) and 639 or 1000 transcripts were down-regulated in response to dbcAMP or 8-CPT-cAMP, respectively. In comparison, the response to butyrate involved many genes: 2091 transcripts were up-regulated in response to butyrate, and 2383 transcripts were down-regulated.

**Figure 2.**
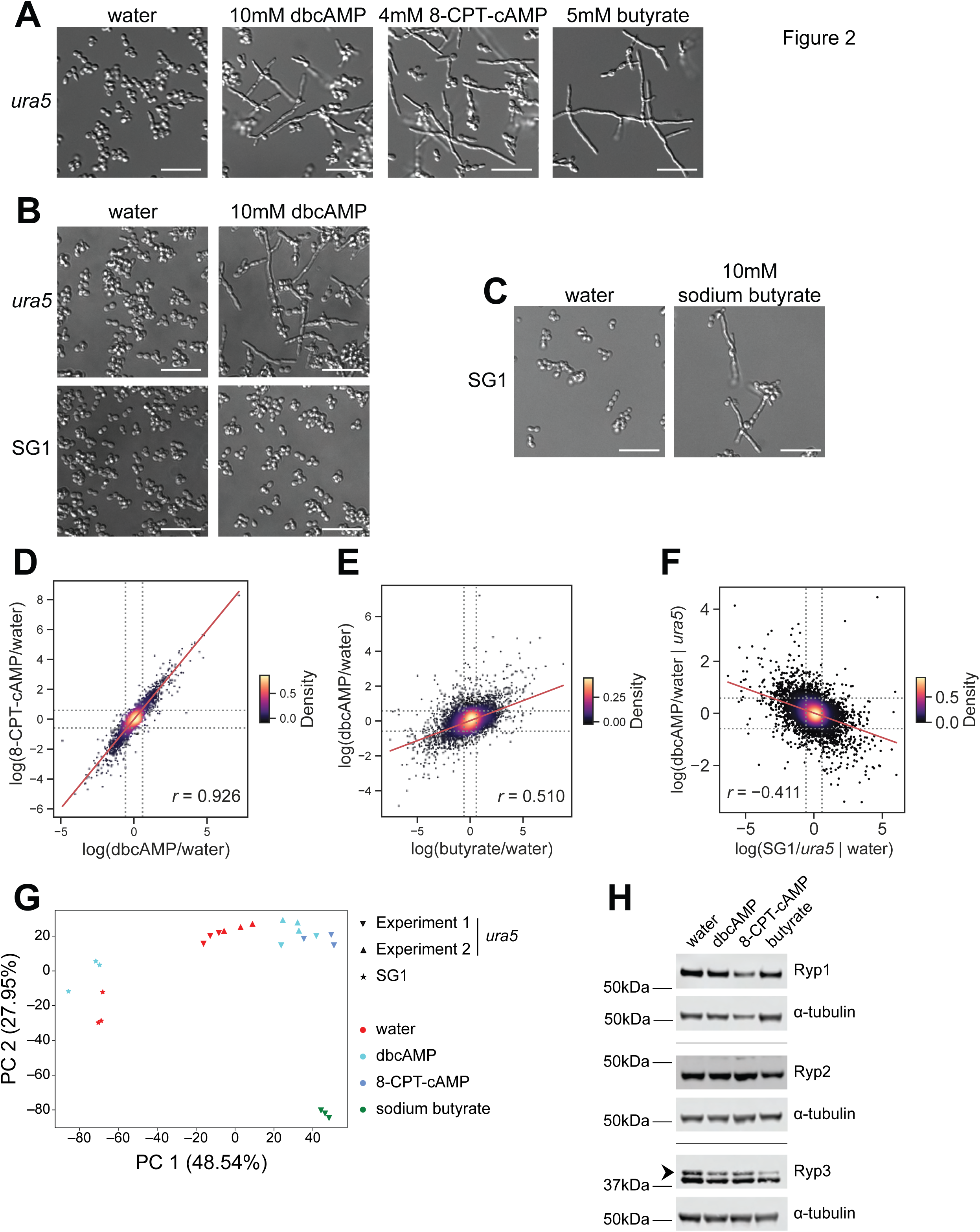
The transcriptional response to cAMP analogs and butyrate. (A) Cell morphology of G217B*ura5 Histoplasma* after 2 days of growth at 37°C in liquid HMM containing GlcNAc with 10mM dbcAMP, 4mM 8-CPT-cAMP, 5mM butyrate, or water as vehicle control. (B) Cell morphology of G217B*ura5* and SG1 *Histoplasma* after 2 days of growth at 37°C in liquid HMM containing GlcNAc with 10mM dbcAMP or water as vehicle control. (C) Cell morphology of SG1 *Histoplasma* after 2 days of growth at 37°C in liquid HMM containing GlcNAc with 10mM sodium butyrate or water as vehicle control. (D, E) Scatter plots of log_2_ LIMMA-fit treatment/vehicle contrasts; dashed line indicates fold-change of log_2_1.5. (F) Scatter plots of log_2_ LIMMA-fit contrasts for genotype (with vehicle) versus treatment (in G217B*ura5*); dashed line indicates fold-change of log_2_1.5. (G) Principal component analysis (PCA) of all genes expressed in G217B*ura5* and SG1 *Histoplasma* treated with dbcAMP, 8-CPT-cAMP, sodium butyrate, or water. In parentheses: percent of total variance explained by the principal component. (H) Western blots performed on protein extracted from G217B*ura5 Histoplasma* after 2 days of growth at 37°C in liquid HMM containing GlcNAc with 10mM dbcAMP, 4mM 8-CPT-cAMP, 5mM butyrate, or water as vehicle control; arrow distinguishes Ryp3 band from known background band. Scale bar denotes 10 µm; *r*, Pearson’s correlation coefficient.

A previous forward genetic screen for yeast-locked mutants identified SG1, a WU15-derived strain that is refractory to filamentous growth in response to acute room temperature shift (11). In contrast to wild-type cells, SG1 did not filament in response to 10 mM dbcAMP at 37°C (Fig. 2B). However, SG1 did filament in response to 10 mM butyrate, albeit to a lesser extent than the wild-type strain (Fig. 2C), highlighting the differential effects of dbcAMP and butyrate addition. Given the distinct morphological response of SG1 to dbcAMP compared to wild-type *Histoplasma*, we performed a second transcriptional profiling experiment comparing the responses of G217B*ura5* and SG1 yeast to dbcAMP treatment.

The effects of cAMP analogs on the transcriptome of G217B*ura5* are highly correlated (Fig 2D, Pearson’s *r* = 0.9). In contrast, the transcriptional response to butyrate is overlapping but distinct (Fig 2E, *r* = 0.5). The transcriptional changes due to the SG1 genotype are weakly anti-correlated to the effect of dbcAMP on G217B*ura5* (Fig 2F, *r* = −0.4), indicating that SG1 responds differently than G217B*ura5* to cAMP analogs transcriptionally as well as morphologically. Simultaneously considering the correlations among all of the transcriptional profiles with principal components analysis (PCA) reveals a first component, representing half of the total variance, that captures the correlated response to dbcAMP and butyrate, anticorrelated with the effects of the SG1 genotype, and a second component, representing half of the remaining variance, capturing an orthogonal response unique to butyrate (Fig 2G).

To stringently define high confidence regulons specific to yeast or hyphal morphology, we progressively gated the transcriptome on expression values that change when the cells are given a filamentation signal in a manner that is dependent on genotype (wild-type or SG1) (Fig. S2). 1270 transcripts were up-regulated (i.e. showed increased abundance) in response to either cAMP analog. Of these, 594 were also up-regulated in response to butyrate. Of these, 436 were also induced at RT in at least one of two previously published room temperature experiments. Of these, 207 were down-regulated (i.e. showed decreased abundance) in the non-filamenting SG1 mutant. Finally, 117 of these transcripts were also down in response to dbcAMP in the SG1 experiment, and we designated this set as the stringently-defined hyphal-associated set (Class 1 in Fig. 3A, B). Likewise, 1186 transcripts were down regulated in response to either cAMP analog. Of these, 842 were also down regulated in response to butyrate. Of these, 563 were also repressed at RT in at least one of the previously published room temperature experiments. Of these, 265 were up-regulated in the non-filamenting SG1 mutant. Finally, a high confidence yeast associated subset of 97 of these genes were also up in response to dbcAMP in the SG1 experiment. We designated this set as the stringently defined yeast-associated set (Class 12 in Fig. 3A, C).

**Figure 3.**
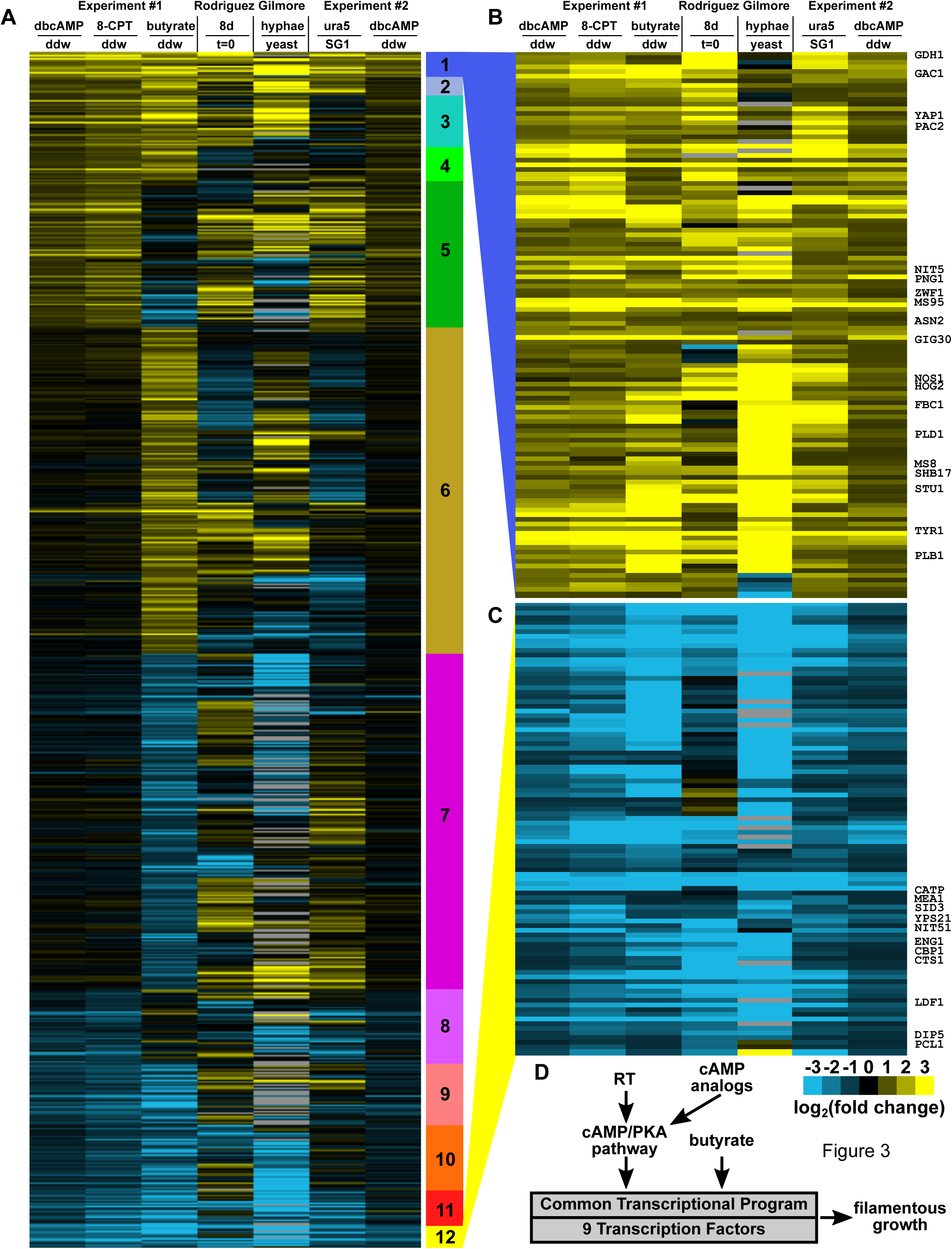
Genetic and environmental perturbations define high confidence morphology regulons. (A) Heatmap showing 5494 genes differentially expressed in response to butyrate, dbcAMP, or 8-CPT-cAMP (Table S3). Genes are grouped by expression pattern according to the scheme in Fig. S2 as indicated by the colored bar to the right of the heatmap. Heatmap colors indicate log_2_ ratios of cpm values. (B) Expanded view of the high confidence yeast regulon (class 1, 112 genes, Table S5). (C) Expanded view of the high confidence hyphal regulon (class 12, 93 genes, Table S6). (D) Summary of experiments and analysis used to identify 9 TFs that increase in abundance under multiple filamentation conditions.

The stringently defined yeast associated set includes the known virulence factors *CBP1* (49–52), *CATP* (53, 54), *ENG1* (55), *LDF1* (50), and the siderophore biosynthetic genes *SID3* and *SID4* (56). The siderophore biosynthetic cluster as a whole is more down regulated in cAMP than in butyrate, with the remaining genes in the cluster passing our significance criteria only for cAMP. More broadly, genes down regulated in response to butyrate, cAMP, and RT include the additional virulence factor *SOD3* (57); regulator of bud site selection *RSR1*; yeast cell wall associated enzymes *AMY3* (58) and *CTS1*/*CHS1*; and GH17/*CFP4*, a secreted yeast gene of unknown function (59, 60). The ER chaperones *KAR2* (BiP), *LHS1*, and *CNE1* (calnexin) are likewise downregulated, as is *OCH1*, an initiator of N-linked glycosylation. This group is also enriched for genes whose promotors are bound by Ryp1 (p = 7.5e-3), Ryp2 (p = 1.7e-4), and Ryp3 (p = 7.5e-4, Benjamini-Hochberg corrected hypergeometric test), the transcription factors that drive yeast-phase growth.

In contrast, the stringently defined hyphal-associated set (Class 1 in Fig. 3A, B) includes anabolic enzymes associated with lipid and riboneogenesis, previously identified hyphal-specific factors, and a variety of signaling molecules of which nine are transcription factors. Zamith-Miranda et al have annotated 77 lipid metabolic genes in Histoplasma capsulatum G186AR (61), 71 of which have G217B orthologs with detectable transcripts in our data set. These genes are significantly up-regulated in butyrate vs. water (p = 1.3e-5) and to a lesser extent in cAMP vs. water (p = 4.9e-4, Wilcoxon rank-sum test). In particular, we do not observe any of the lipid metabolic genes down regulated in response to dbcAMP, only one down in response to 8-CPT-cAMP, and only three down in response to butyrate. Lipid metabolic genes upregulated only in butyrate include the ergosterol pathway. Lipid metabolic genes upregulated in class 1 include the core fatty acid synthase (FAS1 and FAS2) as well as an elongase (ELO2/GIG30). It may be that increased lipid production modulates hyphal membrane fluidity at RT and thus is coupled to the hyphal program even when filamentation is inappropriately induced at 37°C by chemical stimuli.

The reducing potential to drive fatty acid biosynthesis can be derived from isocitrate dehydrogenase or the oxidative arm of the pentose phosphate pathway (62). While isocitrate dehydrogenase (*IDH1*) is not differential in our dataset, *ZWF1*, which catalyzes the first committed step of the oxidative pentose phosphate pathway, yielding NADPH, is in class 1. *GND1*, which catalyzes the other NADPH generating step of this pathway, is upregulated in butyrate but not cAMP. Intriguingly, *SHB17*, which catalyzes the first committed step in the non-oxidative arm of the pentose phosphate pathway is also in class 1. Therefore, these conditions are generating both reducing potential to drive fatty acid biosynthesis, via *ZWF1*, as well as riboneogenesis via *SHB17*, possibly due to increased protein synthesis to replace yeast-specific proteins with hyphal-specific proteins during the transition. While we do not detect upregulation of translational machinery in class 1, GO enrichment analysis reveals that 200 ribosome, pre-ribosome, or rRNA processing genes, as well as 28 tRNA processing genes are upregulated in butyrate (class 6). This set of 228, which is down by a median of 2.4-fold in butyrate vs. water, is also subtly down in 8-CPT-cAMP vs. water (9%) and dbcAMP (13%) and significantly distinguishable from the remainder of the transcriptome (p < 1e-4 in both cases, Wilcoxon rank sum test), suggesting a greater degree of protein turnover corresponding to the stronger hyphal induction in butyrate treated cells. In addition to its roles in lipid biosynthesis and riboneogenesis, we note that *Saccharomyces* null mutants of *ZWF1* require an organic sulfur source to grow (63). Therefore, our observed hyphal-specific expression of *ZWF1* is interesting given the known cysteine auxotrophy of *Histoplasma* yeast but not hyphae (64). On a related note, we see upregulation of the sulfur assimilation pathway in response to filamentation: the first and last genes (*SUL1* and *MET17A*) are in class 2, another (*MET10*) is up in butyrate, and the remainder (*MET3*, *MET14*, *MET16*, and *MET5*) are up in butyrate vs. water, cAMP vs. water, and RT/37°C but not down in SG1/G217B*ura5* (class 3). Additionally, 15 genes involved in amino acid synthesis pathways are upregulated in response to dbcAMP.

In additional to the annotated lipid genes (61), we note three distinct phospholipases in class 1. One phospholipase B (*PLB1*) and one phospholipase D (*PDL1*) have plausible roles in signaling. In particular, the *PLD1* homolog *SPO14* has regulatory roles in meiosis and sporulation in Saccharomyces. Edwards et al previously noted higher expression of *PLB1* and *PLD1* in G186AR vs. G217B yeast (5). The third phospholipase, *PLD2*, which has a conserved secretion signal sequence, is from a distinct lineage of phospholipase D toxins; consistent with toxin function, the *Coccidioides* ortholog of *PLD2* cyclizes, rather than cleaves, lipids (65).

A number of regulatory genes are in class 1, including protein kinases *NIT5* and *HOG2*, protein phosphatase *GAC1*, and the G protein coupled receptor *gprM*. In addition, there are nine transcription factors: *ZNC1*, *YAP1*, *sclB*, *nosA*, *STU1*, *PAC2*, *FBC1*, and two TFs of unknown function (ucsf_hc.01_1.G217B.00081 and ucsf_hc.01_1.G217B.09028). In the dimorphic fungus *Yarrowia lipolytica*, Znc1 is a negative regulator of hyphal growth (66), and sclB is a known developmental regulator in Aspergillus. Of 408 differential genes from RNAseq of *sclB^-^*/WT (67), 31 sclB-repressed and 68 sclB-induced genes have *Histoplasma* orthologs observed in our data. Class 1 contains none of the repressed and five of the induced genes; *viz*., the TFs *FBC1* and ucsf_hc.01_1.G217B.00081 and three enzymes: ucsf_hc.01_1.G217B.03634, ucsf_hc.01_1.G217B.01297, and ucsf_hc.01_1.G217B.08874. Orthologs of *nosA*, *FBC1* and *STU1* are likewise known developmental regulators in Aspergillus (68–70), and in Histoplasma *STU1* overexpression is sufficient for hyphal growth at 37°C (11).

To determine if cAMP analog/butyrate addition affects known regulators of yeast-phase growth, we examined our transcriptional profiles for levels of *RYP1-4* transcripts. *RYP1* and *RYP4* transcript levels were not affected by cAMP analogs but were reduced in butyrate-treated samples; *RYP2* transcript levels were reduced in response to cAMP analogs only in one of the two experiments; and *RYP3* levels were not affected by any treatments. Since Ryp1 and Ryp2 are known to be post-transcriptionally regulated (3), we reasoned that cAMP and/or butyrate could affect Ryp protein levels. We performed Western blot analysis to assay the levels of the Ryp1, Ryp2, and Ryp3 proteins in cultures grown in the presence of dbcAMP, 8-CPT-cAMP, or butyrate. Ryp1 and Ryp2 protein levels appeared to be unchanged in response to these chemicals. In contrast, Ryp3 levels were reduced in response to dbcAMP and 8-CPT-cAMP and further reduced in response to butyrate (Fig. 2H). These observations suggest that a reduction in Ryp3 levels may contribute to filamentation in response to these chemical perturbations.

### The WOPR TF *PAC2* and the C2H2 TF *FBC1* are necessary for cAMP-induced filamentation

Our transcriptional analysis indicated that 9 TFs change in expression in our dataset of interest (Fig. S3 and 3D). We chose to focus on three of these TFs, all of which have orthologs that affect morphology and basic biology of other fungi. We chose the APSES TF *STU1*, which has previously been shown to affect aerial hyphae formation in *Histoplasma* (71) and whose ectopic expression stimulates inappropriate filamentation at 37°C (11). *STU1* is homologous to the *Candida albicans* TF *EFG1*. In *C. albicans*, cAMP production in response to diverse filamentation cues has been shown to activate protein kinase A, in turn activating Efg1 to promote filamentous growth and repress the transcriptional program governed by TFs such as the WOPR TF White-opaque regulator 1 (Wor1) (30). In addition, we subjected the WOPR TF *PAC2*, whose ortholog in *Schizosaccharomyces pombe* has a role in repression of mating (72), to further study. We also examined the role of *FBC1*, a C2H2 transcription factor orthologous to *Aspergillus* spp. FlbC, which is necessary for appropriate conidiation (70).

To test whether *STU1*, *FBC1*, or *PAC2* are necessary for filamentation in response to exogenous cAMP or in response to room temperature, we used CRISPR/Cas9 technology (73, 74) to delete the open reading frame of these genes, or to disrupt them by introducing an indel, causing a frame shift and premature termination (Fig. 4A). The *stu1* mutant exhibited filamentous growth in response to dbcAMP or to room temperature, much like the parental wild-type strain (Fig. 4B and 4C), indicating that *STU1* is not necessary for dbcAMP- or room temperature-induced filamentation in liquid media. In contrast, neither the *pac2* nor the *fbc1* mutant produced hyphae when exposed to dbcAMP, indicating that dbcAMP-induced filamentation is dependent on each of these TFs. Interestingly, both mutants were able to transition to hyphal growth within 8 days of room temperature growth, although under these conditions there is less filamentation in the *fbc1* mutant strain (Fig. 4C). Nonetheless, *PAC2* and *FBC1* are individually dispensable for room temperature-induced filamentation.

**Figure 4.**
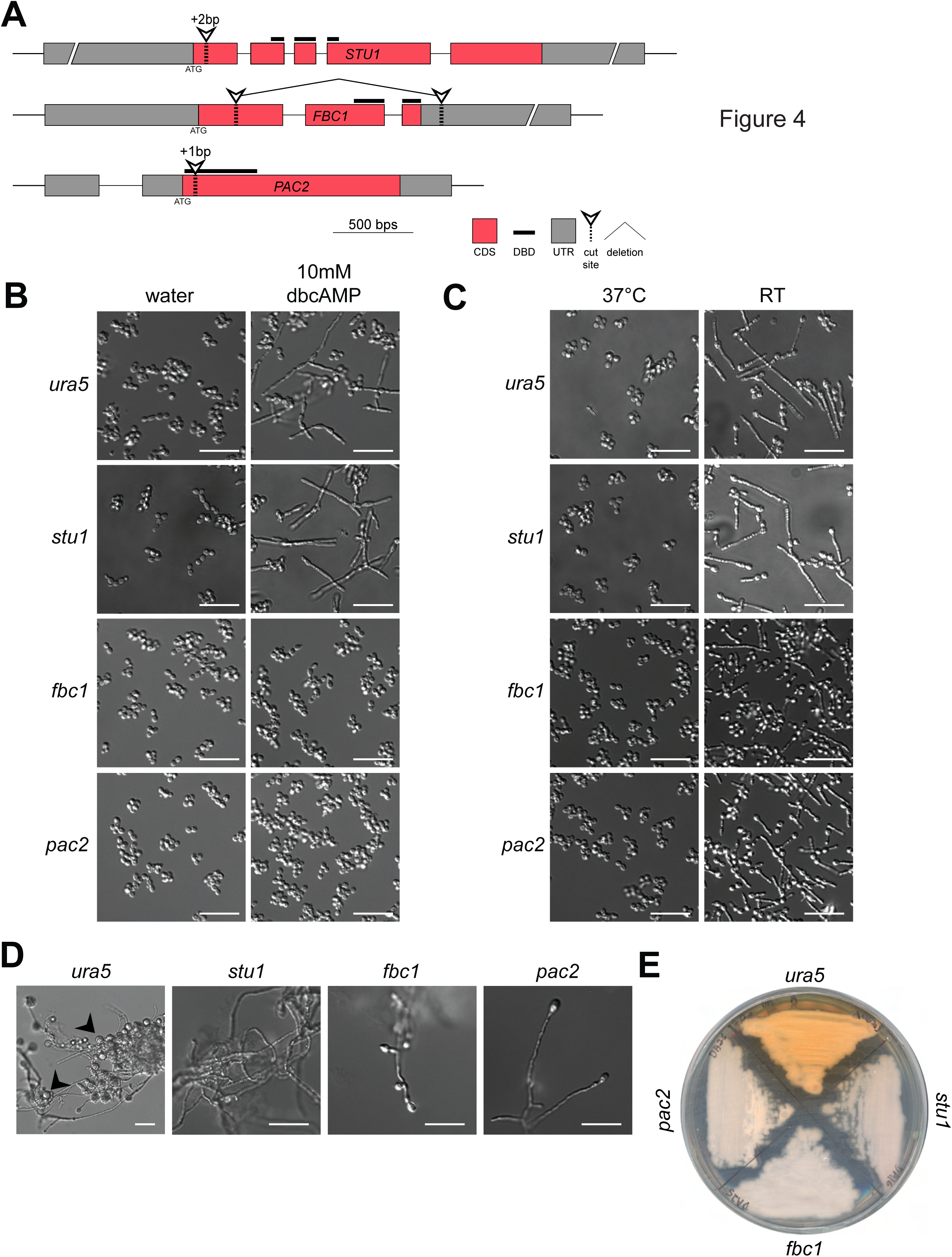
Transcription factors *PAC2* and *FBC1* are necessary for cAMP-induced filamentation. (A) Scheme summarizing genetic manipulations of *Histoplasma* using CRISPR/Cas9 to disrupt *STU1* and *PAC2*, and delete the open reading frame of *FBC1*. (B) Cell morphology of G217B*ura5* (parental strain), *stu1*, *pac2*, and *fbc1 Histoplasma* after 2 days of growth at 37°C in liquid HMM containing GlcNAc with or without 10mM dbcAMP. (C) Cell morphology of G217B*ura5*, *stu1*, *pac2*, and *fbc1 Histoplasma* after 8 days of growth at RT in liquid HMM containing GlcNAc. (D) Cell morphology of G217B*ura5* (parental strain), *stu1*, *pac2*, and *fbc1 Histoplasma* after 8 days of growth at RT on solid Sabouraud media. Arrows indicate macroconidia. (E) Gross growth morphology of G217B*ura5* (parental strain), *stu1*, *pac2*, and *fbc1 Histoplasma* after 14 days of growth at RT on solid HMM+GlcNAc media. bp: base-pair; CDS: coding sequence; DBD: DNA binding domain; UTR: untranslated region; scale bar denotes 10 µm.

Since fungi exhibit morphologic changes on solid medium, we examined the phenotype of G217B*ura5* (parental strain), *stu1*, *pac2*, and *fbc1* strains on solid Sabouraud media at room temperature. We observed that the parental strain generated large amounts of macroconidia (Fig. 4D). In contrast, the *stu1*, *pac2*, and *fbc1* mutants were each deficient in the formation of these cells (Fig. 4D). Additionally, when grown on HMM plates containing GlcNAc at room temperature, these mutants yielded glossy, pink colonies as opposed to the fuzzy orange-brown colonies of the parental G217B*ura5* (Fig. 4E), although both colonies consisted of filamentous cells (data not shown). Taken together, these data indicate that each of these transcription factors is required for the normal developmental program, including macroconidia formation and pigment production, on solid medium at room temperature.

### Ectopic expression of *PAC2* and *FBC1* stimulates inappropriate filamentation at 37°C

Ectopic expression of *STU1* is sufficient to override the yeast program and promote robust filamentation at 37°C (11). To test whether deregulated expression of *PAC2* and *FBC1* has a similar phenotype at 37°C, we used a constitutive *ACT1* promoter to drive their expression. Ectopic expression of *PAC2* resulted in filamentous colonies at 37°C, while ectopic expression of *FBC1* resulted in an aberrant wrinkled colony at 37°C containing both yeast and hyphae (Fig. 5A). To understand the relationships between the activity of the three transcription factors, we also ectopically expressed each one of them in the background of the three mutants. Overexpression of *FBC1* in the *stu1* mutant did not result in filamentation at 37°C, and overexpression of *PAC2* in the *stu1* mutant resulted in a mixed yeast-hyphae morphology. In contrast, ectopic expression of *STU1* triggered filamentation at 37°C in both the *fbc1* and *pac2* mutants, suggesting that Stu1 functions downstream of Fbc1 and Pac2. In contrast, ectopic expression of *FBC1* had a much less dramatic phenotype in the *pac2* mutant compared to the parental strain, indicating that the effect of *FBC1* overexpression is dependent on *PAC2*. Conversely, ectopic expression of *PAC2* was able to promote filamentous growth at 37°C in both the *fbc1* and *pac2* mutants (Fig. 5A). Taken together, these results show that the ability of Fbc1 to induce filamentous growth at 37°C depends on the activity of Pac2 and Stu1, and similarly Fbc1- and Pac2-dependent filamentation at 37°C depends on Stu1 (Fig. 5B).

**Figure 5.**
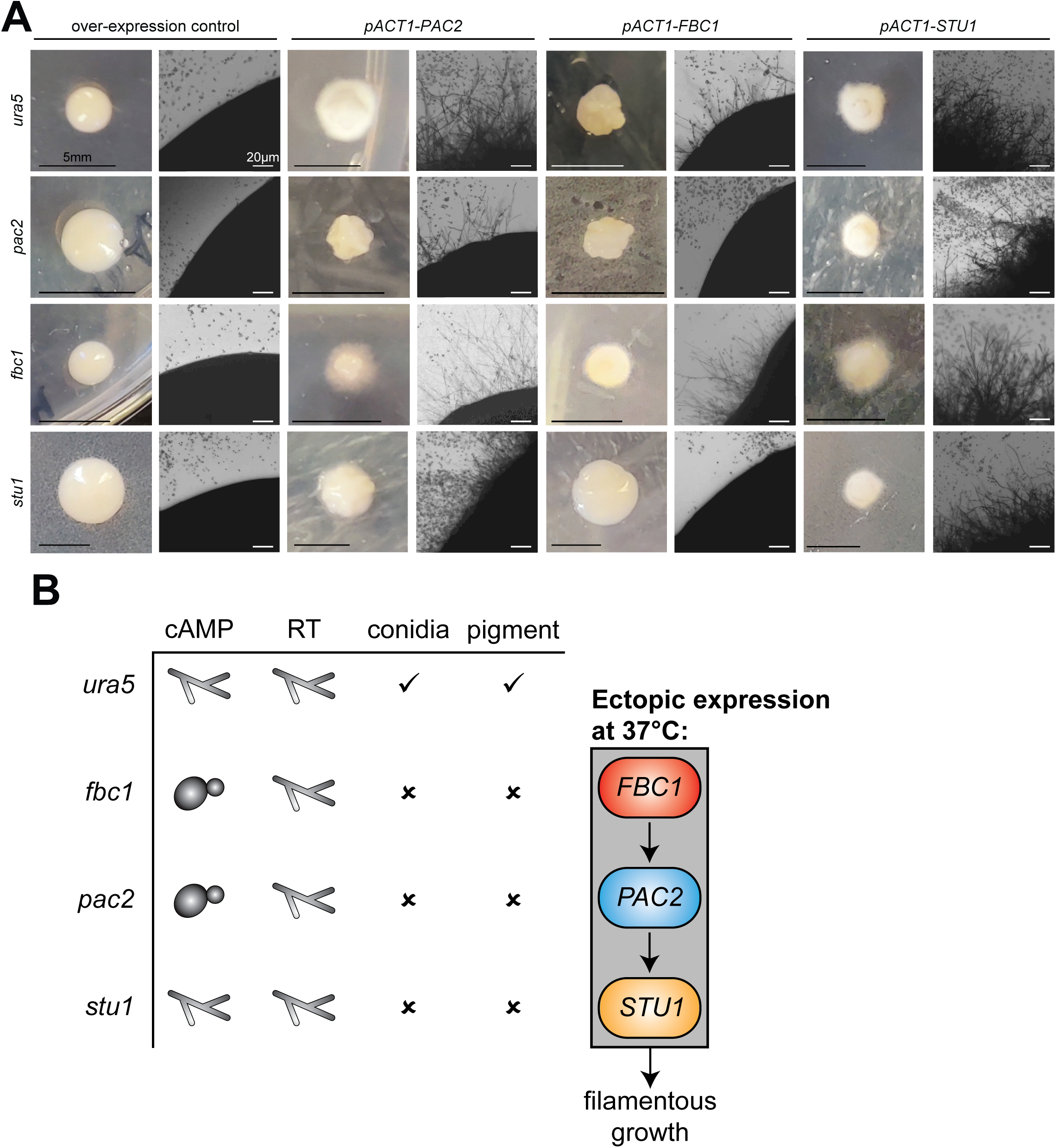
Expression of *PAC2* and *FBC1* is sufficient to trigger inappropriate filamentation at 37°C. (A) Various *Histoplasma* strains (genotype indicated on the left) were transformed with ectopic expression constructs carrying *PAC2*, *FBC1*, or *STU1* (indicated on the top). For each transformation, a macroscopic image of colonies on the transformation plate is shown on the left, and a microscopic image of the corresponding colony edge is shown on the right. Untransformed microcolonies can be seen as small dark dots in the background of the microscopic images. (B) Table summarizing the *fbc1*, *pac2*, and *stu1* phenotypes and a model showing the genetic relationship between *FBC1*, *PAC2*, and *STU1* in the context of ectopic expression at 37°C.

## Discussion

In this study we have shown that exogenous cAMP is sufficient to promote filamentous growth in *Histoplasma* by triggering transcriptional reprogramming at 37°C. We compared the transcriptional response to three filamentation signals—the classic physiological signal of RT, exogenous cAMP at 37°C, and exogenous butyrate at 37°C. We determined that 9 transcription factors are induced by all three of these signals and hypothesized that these factors regulate the hyphal growth program. We focused on three such factors that were induced by cAMP and repressed in the non-filamenting mutant SG1: *STU1*, *FBC1*, and *PAC2*. We have shown that while *STU1* is dispensable for cAMP-induced filamentation, *PAC2* and *FBC1* each are necessary for cAMP-induced filamentation. Each of these transcription factors is not individually necessary for room temperature-induced filamentation in liquid media, although each transcription factor is necessary for normal hyphal development (including macroconidia and pigment production) on solid media. Finally, we have shown that ectopic expression of *STU1*, *FBC1*, or *PAC2* is sufficient to drive filamentous growth at 37°C, that the ability of *FBC1* to promote filamentous growth at 37°C is dependent on *PAC2*, and that the ability of *FBC1* and *PAC2* to promote filamentous growth depends on *STU1*, suggesting the transcription factor hierarchy depicted in Figure 5B.

We utilized two chemical stimuli that promote filamentation despite growth at 37°C: cyclic AMP and butyrate. While the ability of exogenous cAMP to promote filamentation in *Histoplasma* has been described in the literature (47), the transcriptional changes that occur during cAMP treatment were unknown. Additionally, this study is the first to observe the ability of butyrate to promote filamentous growth. We initially investigated butyrate to determine if its release from dibutyryl-cAMP, a cAMP analog commonly used to study PKA activation, was responsible for the response of *Histoplasma* to dbcAMP. However, the response of *Histoplasma* to butyrate is distinct from its response to dbcAMP: Butyrate is a much more potent filamentation stimulus, it triggers a unique transcriptional response, and the transcriptional response of *Histoplasma* to dbcAMP is near-identical to its transcriptional response to 8-CPT-cAMP, which does not contain butyrate. Thus, we conclude that the effects of dbcAMP are not due to release of butyrate from this compound but rather via its activity in the cAMP pathway.

To identify gene sets that correspond with the morphological transition to filamentous growth, we conducted expression profiling of cultures filamenting in response to dbcAMP and to butyrate. About 46% of the total transcriptome is differentially expressed in samples treated with butyrate. Specifically, we detected up-regulation of lipid metabolic genes, ergosterol synthesis genes, and ribosomal proteins in response to butyrate. While the up-regulation of ribosomal proteins may be connected with the change in the proteome that is necessary to establish a different growth form (yeast vs. filaments), the role of lipid metabolism genes in this context remains to be elucidated.

It is unclear whether the transcriptional response to butyrate reflects a physiologically relevant response to butyrate as a filamentation inducing signal; e.g., *Histoplasma* may naturally encounter soil-residing bacteria, which are capable of producing butyrate (75). Interestingly, butyrate can act as a pan-histone deacetylase inhibitor (HDACi) (76). In other fungi, HDACi (including butyrate) have been shown to affect developmental programs: a study conducted in *C. albicans*, *C. parapsilosis*, and *C. neoformans* found that butyrate decreased fungal growth and biofilm formation in *Candida albicans* and *C. parapsilosis*, inhibited filamentation in *C. albicans*, and prevented capsule formation and melanization in *C. neoformans* (77). Introduction of butyrate or other HDACi, as well as genetic disruption of HDACs, has also been shown to modulate diverse processes such as virulence, morphology, conidiation, germination, and the expression of secondary metabolite (SM) gene clusters in *Aspergillus*, *Magnaporthe*, *Cryptococcus*, *Cochiobolus carbonum*, and others (78–83).

In contrast to butyrate, the cAMP analogs dbcAMP and 8-CPT-cAMP only affected the expression of 25% of the transcriptome. We did not detect any concerted regulation of carbon utilization pathways in our data in response to cAMP. However, 15 genes involved in amino acid synthesis pathways are upregulated in response to dbcAMP, as well as both subunits of the fatty acid synthase (FAS) complex, suggesting that, similar to *S. cerevisiae* (16, 84, 85), *Histoplasma* up-regulates biosynthetic pathways in response to cAMP. Interestingly, the OLE1 Δ9-desaturase, which Storlazzi *et al*. have previously reported to be down-regulated in response to exogenous cAMP (86), is up-regulated in response to cAMP in our experiments. *OLE1* is not differentially expressed in the steady-state yeast or mycelial phases of *Histoplasma* G217B (3), but appears to be consistently upregulated in the days following the transition of *Histoplasma* cultures to room temperature (11). These data are contrary to the finding reported by Gargano *et al.* (87) that *OLE1* transcript was present in G217B yeast but absent in mycelia. Future experiments to target *OLE1* levels could address the function of this gene in morphogenesis. For example, increase of Ole1 activity could lead to elevated levels of unsaturated fatty acids in cellular membranes, with potential consequences on signaling cascades, thermotolerance and virulence.

By comparing differential expression induced by both cAMP analogs and butyrate with existing data for differential expression between 37°C and RT and between WT *Histoplasma* and the non-filamenting SG1 mutant, we arrived at high-confidence filamentation- and yeast-associated gene groups. By definition, these groups are well-correlated with previously defined morphology-associated groups that compared gene expression in response to room temperature with that of SG1 (11). The stringent yeast group is enriched for Ryp1, Ryp2 and Ryp3 direct targets, and contains several genes encoding small, secreted proteins and genes involved in iron acquisition, all hallmarks of the *Histoplasma* yeast phase. Interestingly, this group is largely devoid of genes encoding known regulators and transcription factors. An exception is Mea1, the ortholog of the *A. nidulans* nitrogen regulator MeaB, which is involved in the regulation of nitrogen metabolism (88).

In contrast, the stringent gene group associated with filamentous growth contains multiple transcripts encoding signaling molecules and transcription factors, as well as transcripts that suggest an upregulation of the pentose phosphate pathway (*ZWF1*, *SHB17*), lipid biosynthesis (*FAS1*, *FAS2*, *GIG30*), and lipid processing (phospholipases B and D). A connection between cell morphology and the metabolic state of *Histoplasma* remains to be elucidated in future studies.

The stringent hyphal-associated gene group contains 9 genes predicted to function as transcription factors, of which 6 have previously been associated with regulation of morphology and development in other fungi: the Znc1 ortholog in *Y. lipolytica* regulates filamentation (66); sclB regulates conidiation and SM production in *A. nidulans* (67); *STU1* encodes an APSES transcription factor, and its orthologs stuA and Efg1 regulate development and morphology in *Aspergillus* and *Candida*, respectively (15, 35, 69); *FBC1* ortholog *flbC* regulates conidiation in *Aspergillus* (70); *NosA* regulates sexual development in *A. nidulans* (89); and *PAC2* encodes a WOPR family transcription factor, orthologous to S. pombe *pac-2*, which represses the cAMP-dependent expression of *ste-11*, thus inhibiting mating (72). Together, this group of transcription factors are attractive candidates for regulators that are relevant to the ability of *Histoplasma* to switch between the yeast and hyphal morphology.

We chose to further characterize the role of three of these transcription factors in promoting filamentation in response to cAMP and RT: *PAC2*, *FBC1*, and *STU1*. While *PAC2* and *FBC1* were necessary to trigger filamentous growth in response to cAMP, *STU1* was dispensable for cAMP-dependent filamentation, and none of these three transcription factors were necessary for RT-induced filamentation in liquid culture. Although we reported previously that disruption in the promoter region of *STU1* has a partial filamentation phenotype, we did not observe the same phenotype in liquid culture in the ORF-disruption mutant we generated in this study. However, we did observe a RT phenotype on plates. In fact, mutants lacking functional Pac2, Fbc1, or Stu1 each had developmental defects in the full hyphal program at RT (Fig. 4D, E). Additionally, we found that overexpression of each of the transcription factors was sufficient to drive filamentous growth at 37°C (Fig. 5A). Additional work is needed to determine whether these, and other transcription factors whose expression is correlated with filamentous growth, act redundantly to drive filamentation in liquid culture in response to temperature.

Our ectopic expression work uncovered interesting relationships between Pac2, Fbc1, and Stu1. We found that *PAC2* was necessary for *FBC1*-driven filamentation, and similarly, *STU1* was necessary for *FBC1* and *PAC2* ectopic expression to cause filamentous growth at 37°C. Together, this suggests a genetic pathway with *PAC2* downstream of *FBC1*, and *STU1* downstream of *PAC2* (Fig. 5B). However, regulation of eukaryotic transcription is often complex, and further research is necessary to determine the roles of these 3 transcription factors in controlling and coordinating the transcriptional response to cAMP. We note that *PAC2*, which we have implicated in the hyphal program in this study, and *RYP1*, which is required for the yeast program, are paralogs, suggesting specialization of these individual TFs in opposing developmental programs. Further exploring the circuits that govern the transition between yeast and hyphae in thermally dimorphic fungi will elucidate the critical basic biology of these pathogens and provide an opportunity to develop molecular strategies for targeted therapeutics.

## Acknowledgements

We are grateful to Bruce S. Klein, James B. Konopka, and William E. Goldman for generously sharing reagents and information. We thank Suzanne M. Noble, Alexander D. Johnson, and Jonathan S. Weissman for their guidance. Eric D. Chow and the UCSF Center for Advanced Technology provided invaluable advice on library preparation and sequencing. We also thank members of the Sil and Noble labs for helpful discussions.

## Materials and methods

### *H. capsulatum* strains and growth conditions

All strain manipulations were done in *G217Bura5*-DA, which is derived from the *Histoplasma* G217B*ura5* (WU15) background. Strains used in this paper can be found in Table S1. Strain frozen stocks (8% DMSO or 15% Glycerol) were streaked onto *Histoplasma* Macrophage Medium (HMM) agarose plates supplemented with 0.2mg/ml uracil where necessary. Liquid *Histoplasma* cultures were inoculated from solid media plates into HMM + 100 U/ml Penicillin/ Streptomycin (with 0.2mg/ml of uracil if needed) and were passaged 1:25 every 2-3 days into fresh medium, unless indicated otherwise. The glucose in HMM was replaced with equimolar amounts of N-acetyl-glucosamine (A3286, Sigma Aldrich) in indicated experiments. Where mentioned, dibutyryl cyclic adenosine monophosphate sodium (dbcAMP, D0627, Sigma Aldrich), 8-(4-Chlorophenylthio)-cAMP sodium (8-CPT-cAMP, ab120424, Abcam), 3-isobutyl-1-methylxanthine (IBMX, ab120840, Abcam), sodium butyrate (B5887, Sigma Aldrich), or cyclic adenosine monophosphate sodium (cAMP, A6885, Sigma Aldrich) were added to cultures after dissolving in autoclaved water and filter sterilizing by passing through a 0.22 MCE filter (Santa Cruz Biotechnologies). For 37°C growth, plates were grown in a humidified incubator with 5% CO_2_, and liquid cultures were grown on an orbital shaker at 120-150rpm. For room temperature (RT) growth, plates were wrapped in parafilm and plastic bags and placed in a 25°C incubator in a Biosafety level 3 facility. Liquid RT cultures were grown at ambient temperature (26-28°C) on an orbital shaker at 120 rpm.

### Strains and Cloning

The SG1 mutant was generated using *Agrobacterium*-mediated insertional mutagenesis as described previously (11). Subcloning was performed in *E. coli* DH5α- or DH10b-derived strains. The *STU1* overexpression strain and the control strain were constructed as described previously (11). *FBC1* and *PAC2* overexpression strains were constructed by amplifying the open reading frame (ORF) and 3’ untranslated region (3’UTR) by PCR, and cloned into pTM1 (a pDONR/Zeo based plasmid) downstream of the ACT1 promoter using Gibson cloning kit (NEB), and then introduced into pDG33 using Gateway cloning (Thermo Fisher Scientific). To introduce gene disruption, we used the strategy described previously (28, 29, Joehnk et al, in preparation). Briefly, a gene-appropriate protospacer was inserted into a self-cleaving element using PCR and cloned into the episomal plasmid pBJ219 (which contains the *HcURA5* gene) using Gateway cloning. The plasmid was linearized and introduced into *Histoplasma* G217B*ura5* using electroporation (as described previously in (56)) along with an mCherry-targeting protospacer as a negative control. Transformants that grew on HMM without uracil were assayed for the disruption using PCR amplifying the targeted genomic region followed by sequencing of the product and indel analysis using the ICE tool (inference of CRISPR edits) from Synthego. The colony with the highest indel levels was selected, grown on fresh media, and plated for single colonies. The process was repeated until no wild-type sequence could be detected. The plasmid was then lost by growing the isolate for 3 passages in liquid media containing uracil, plating the culture for single colonies and identifying colonies that were auxotrophic for uracil. To create whole gene deletion mutants, the aforementioned technique was adapted to include two protospacers targeting the sequences 5’ and 3’ of the gene. To test for a deletion, DNA was extracted from a liquid *Histoplasma* culture, and PCR with primers external to the deleted sequence, or with primers internal to the sequence, was performed. The colony with highest amount of editing (short external PCR product, lowest levels of internal PCR product) were selected, grown on fresh media, and plated for single colonies. The process was repeated until no wild-type sequence could be detected.

### Imaging and image analysis

DIC imaging was performed on a Zeiss AxioCam MRM microscope at 40× magnification. Colony images were captured using a Leica microscope. Gross morphology of colonies was captured using a OnePlus 8 Pro camera. To quantify filamentation events, Fiji was used to count yeast and filamentous events in 4 frames per sample.

### Culture conditions for cAMP expression profiling experiments

Samples were prepared in two separate experiments. In the first experiment, *Histoplasma* G217B*ura5* was grown in liquid HMM culture at 37°C for 2 passages. On the day of chemical addition, the culture was passaged into fresh HMM + GlcNAc at a final OD600 of 0.2, and 10 mM dbcAMP, 4 mM 8-CPT-cAMP, 5 mM of sodium butyrate, or filter-sterilized autoclaved water (vehicle control) were added to the appropriate cultures, with 3 biological replicates for each chemical. Cultures were returned to 37°C growth with shaking. 2 days after the addition of each chemical, a small volume of each culture was fixed by adding paraformaldehyde (final paraformaldehyde concentration 4% v/v), and 10 ml of each culture were harvested by filtration using Nalgene Rapid-Flow Sterile Disposable Bottle Top Filters with SFCA Membrane (Fisher 09-740-22G). Cells were scraped from the filter into tubes and promptly flash-frozen in liquid nitrogen.

In the second experiment, *Histoplasma* G217B*ura5* or the SG1 mutant were grown in liquid HMM culture at 37°C for 2 passages. On the day of dbcAMP addition, the cultures were passaged into fresh HMM + GlcNAc at a final OD600 of 0.2, and 10 mM dbcAMP or filter-sterilized autoclaved water (vehicle control) were added to the appropriate cultures, with 3 biological replicates for each chemical-strain combination. The cultures were treated and samples were collected as described for the first experiment.

### RNA extraction

Total RNA was extracted from fungal cells using a Qiazol-based RNA extraction protocol. Frozen cell pellets were resuspended in Qiazol (Qiagen, Germany) and incubated at RT for 5 minutes to thaw. The lysate was subjected to bead beating (Mini-Beadbeater, Biospec Products, Bartlesville, OK) followed by a chloroform extraction. The aqueous phase was then transferred to an Epoch RNA column where the filter was washed with 3 M NaOAc (pH=5.5) and then with 10 mM TrisCl (pH=7.5) in 80% EtOH. DNase (Purelink, Invitrogen, Carlsbad, CA) treatment was used to remove any residual DNA, and the filters were washed again with NaOAc and TrisCl before eluting the RNA in nuclease-free water.

### mRNA isolation

For each experiment, mRNA was extracted from an equal amount of total RNA (up to 20 μg) of each sample. Total RNA samples and were treated with TURBO DNase (Thermo Fisher, Waltham, MA). RNA quality was determined with a RNA 6000 Nano Bioanalyzer chip (Agilent Technologies, Santa Clara, CA). mRNA was purified using polyA selection with Oligo-dT Dynabeads (Thermo Fisher, Waltham, MA) as described in the manufacturer’s protocol. Ribosomal RNA depletion was confirmed with an RNA 6000 Nano Bioanalyzer chip.

### RNAseq library preparation

Libraries for RNAseq were prepared using the NEB Next Ultra II Directional RNA Library Prep Kit (New England Biolabs, Ipswich, MA). Individual libraries were uniquely barcoded with NEBNext Multiplex Oligos for Illumina sequencing platform (New England Biolabs, Ipswich, MA). Average fragment size and presence of excess adapter was determined with High Sensitivity DNA Bioanalyzer chip from Agilent Technologies (Santa Clara, CA). Libraries had an average fragment length of 300 to 500 bp. The concentration of the individual libraries was quantified using the High Sensitivity DNA Qubit assay (Thermo Fisher, Waltham, MA). A total of 5 ng of each library was pooled into each final libraries and run on a High Sensitivity DNA Bioanalyzer chip to determine the average fragment size of the final pooled samples. The final libraries were submitted to the UCSF Center for Advanced Technology for sequencing on an Illumina HiSeq 4000 sequencer.

### Transcriptome analysis

Transcript abundances were quantified based on version ucsf_hc.01_1.G217B of the *Histoplasma* G217B transcriptome (S5 Data of (3)). Relative abundances (reported as TPM values (90)) and estimated counts (est_counts) of each transcript in each sample were estimated by alignment free comparison of k-mers between the reads and mRNA sequences using KALLISTO version 0.46.0 (91). Further analysis was restricted to transcripts with TPM ≥ 10 in at least one sample.

Differentially expressed genes were identified by comparing replicate means for contrasts of interest using LIMMA version 3.30.8 (92, 93). Genes were considered significantly differentially expressed if they were statistically significant (at 5% FDR) with an effect size of at least 1.5x (absolute log2 fold change ≥ 0.585) for a given contrast.

For comparison with previous expression profiling, the above KALLISTO/LIMMA pipeline was applied to the reads from (3) (SRA accession SRP058149) and LIMMA fit parameters were taken from S2 Data of (11). Enrichments for Ryp1-3 targets in gene sets were calculated using a hypergeometric test, followed by Benjamini-Hochberg correction for multiple hypotheses testing, with adjusted p < 0.05 considered significant.

### Protein extraction

Organic fractions from Qiazol-chloroform extraction were stored at –20°C. After thawing the fractions, 100% ethanol was added, and samples were centrifuged to pellet DNA. The protein-containing supernatant was added to isopropanol and centrifuged at 4°C to pellet the protein precipitate. The pellet was washed 3 times with 0.3M guanidinium thiocyanate in 95% ethanol followed by centrifugation at 4°C, after which 100% ethanol was added to the pellets. The protein pellets were vortexed and incubated at RT for 20 minutes, centrifuged at 4°C, and air-dried at RT. The pellets were resuspended in urea lysis buffer (9 M Urea, 25 mM Tris-HCl, 1% SDS, 0.7 M β-mercaptoethanol) and incubated at 50°C up to 20 minutes until fully dissolved in the buffer, followed by boiling and centrifugation. The supernatant was transferred into a clean tube and quantified using the Pierce 660nm assay with added ionic detergent compatibility reagent (Thermo Fisher, Waltham, MA).

### Western Blotting

Following quantification of protein, an equal amount per sample of 12 µg was boiled with Novex NuPAGE LDS Sample Buffer (Invitrogen, Carlsbad, CA) and loaded onto a 10-well Novex NuPAGE 4% to 12% BT SDS-PAGE gel (Invitrogen, Carlsbad, CA). Electrophoresis was performed in MOPS running buffer at 150 V. The protein was then transferred to a nitrocellulose membrane at approximately 35 V for 2 hours. The membrane was incubated with Intercept® (PBS) Blocking Buffer (LI-COR, Lincoln, NE) for an hour and then incubated in the primary antibody in wash buffer overnight at 4 °C. Polyclonal peptide antibodies against either Ryp1, Ryp2, or Ryp3 were used as primary antibodies in the following dilutions in blocking buffer: rabbit anti-Ryp1 (1:10,000), rabbit anti-Ryp2 (1:2,500), rabbit anti-Ryp3 (1:5,000) (4). As a loading control, monoclonal mouse-anti-α-tubulin (1:1,000) was used (DM1A, Novus Biologicals, Littleton, CO). The blot was washed with PBS + 0.1 % Tween-20 three times, and secondary antibody was added to the blot for 1 hour at room temperature: IRDye 800CW Goat anti-Rabbit IgG, or IRDye 680RD Donkey anti-Mouse IgG (both 1:10,000, LI-COR, Lincoln, NE). The blot was washed with PBS + 0.1 % Tween-20 three times, and imaged using the LI-COR Odyssey CLx system.

### *Histoplasma* genomic DNA extraction

Cells from 10 mL of dense liquid culture were collected by centrifugation and kept at –80°C until DNA extraction. The cell pellets were thawed and washed in TE buffer, and then resuspended in lysis buffer (50 mM Tris [pH 7.2], 50 mM EDTA, 0.1 M SDS, 0.14 M β-mercaptoethanol). Cells were lysed in buffer by bead-beating with zirconia beads, and the samples were incubated at 65°C for 1 hour. 800 µl of 1:1 phenol-chloroform solution was added to each sample followed by thorough mixing. The samples were centrifuged for 15 minutes at 13,000 rpm. The aqueous fraction was added to 470 µl of 0.13 M sodium acetate in 95 % isopropanol and mixed by inversion to precipitate the DNA. The supernatant was discarded after a 2-minute centrifugation, and the pellet was washed with 70% ethanol and air-dried at 65°C. 1 mL TE buffer with 1.5 µl RNase A (10mg/ml) was added to each sample and the pellet was dissolved at 65°C.

---

**Figure S1.**
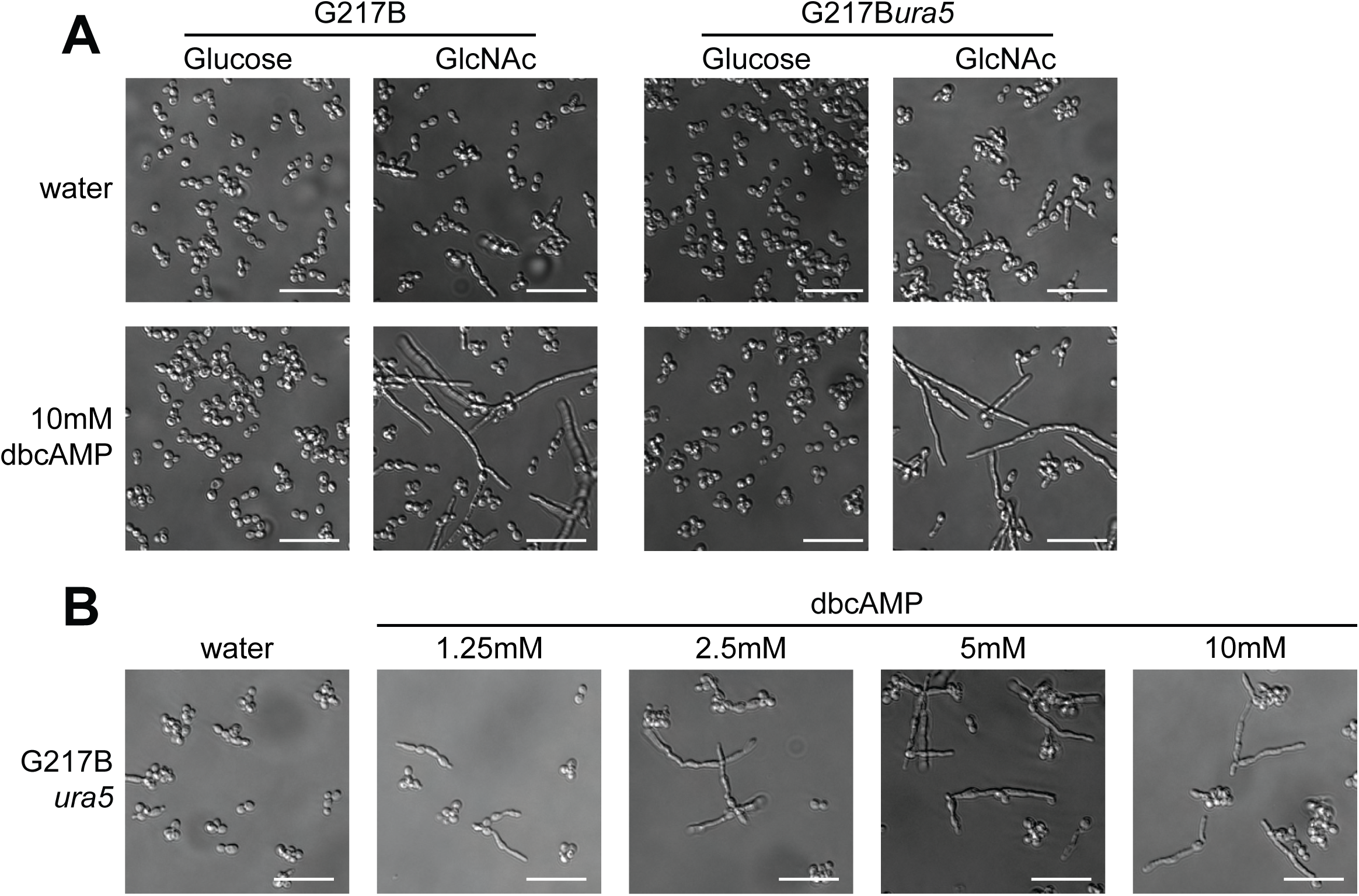
dbcAMP promotes filamentous growth at 37°C. (A) Cell morphology of G217B and G217B*ura5 Histoplasma* after 3 days of growth at 37°C in liquid HMM containing glucose or GlcNAc with or without 10mM dbcAMP. (B) Cell morphology of G217B*ura5 Histoplasma* after 2 days of growth at 37°C in liquid HMM containing GlcNAc with various concentrations of dbcAMP. Scale bar denotes 10 µm.

**Figure S2.**
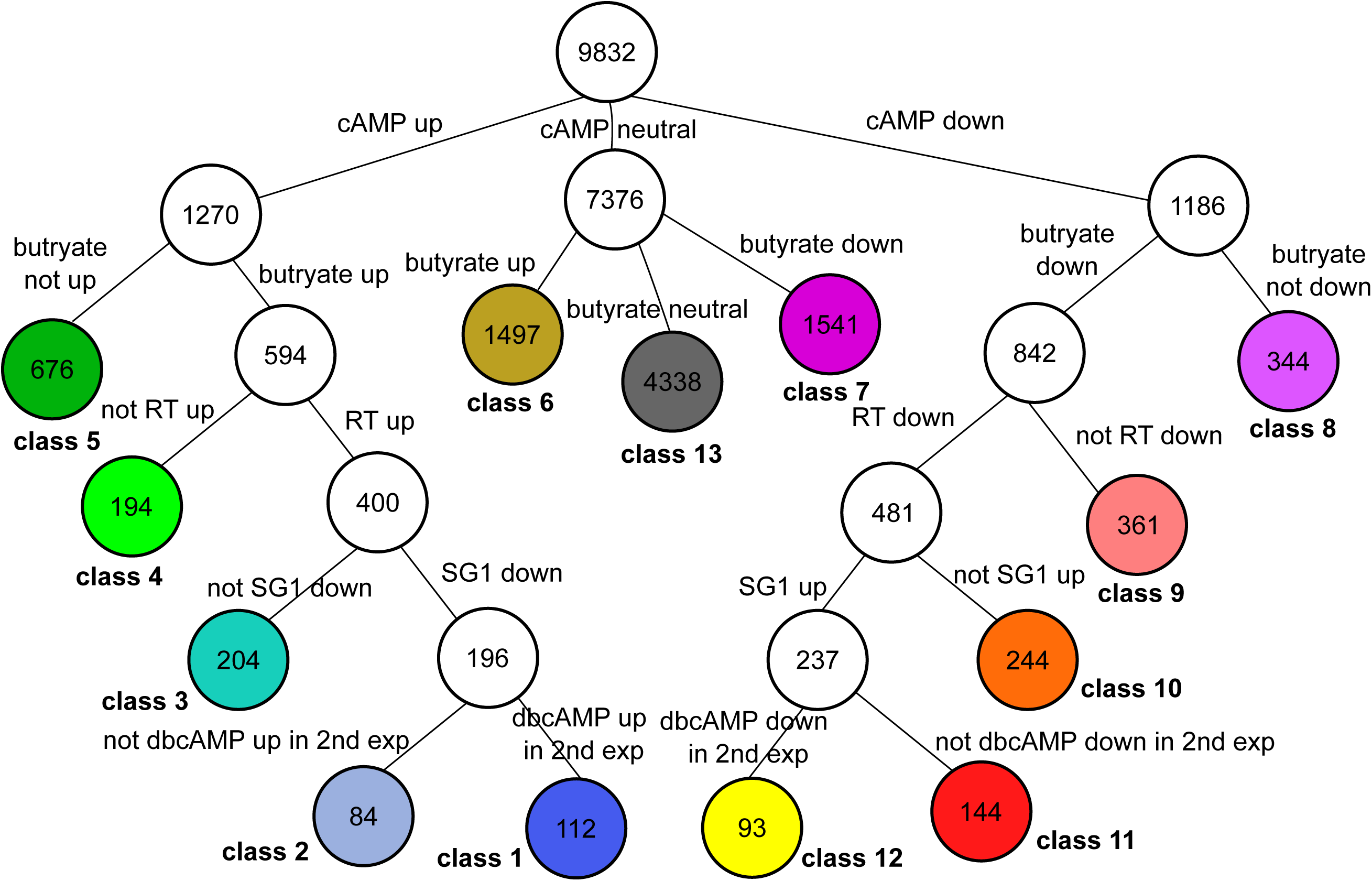
Classification of genes by expression profile. Scheme for the classification of genes as discussed in the text and shown in Fig. 3. The top node indicates the 9832 genes observable in the expression profiling data. Edges divide each internal node into disjoint sets, with edge labels indicating the division criteria. Numbers inside nodes indicate the number of genes in the corresponding set. The colored terminal nodes indicate the final, disjoint classification spanning the 9832 observed genes.

**Figure S3.**
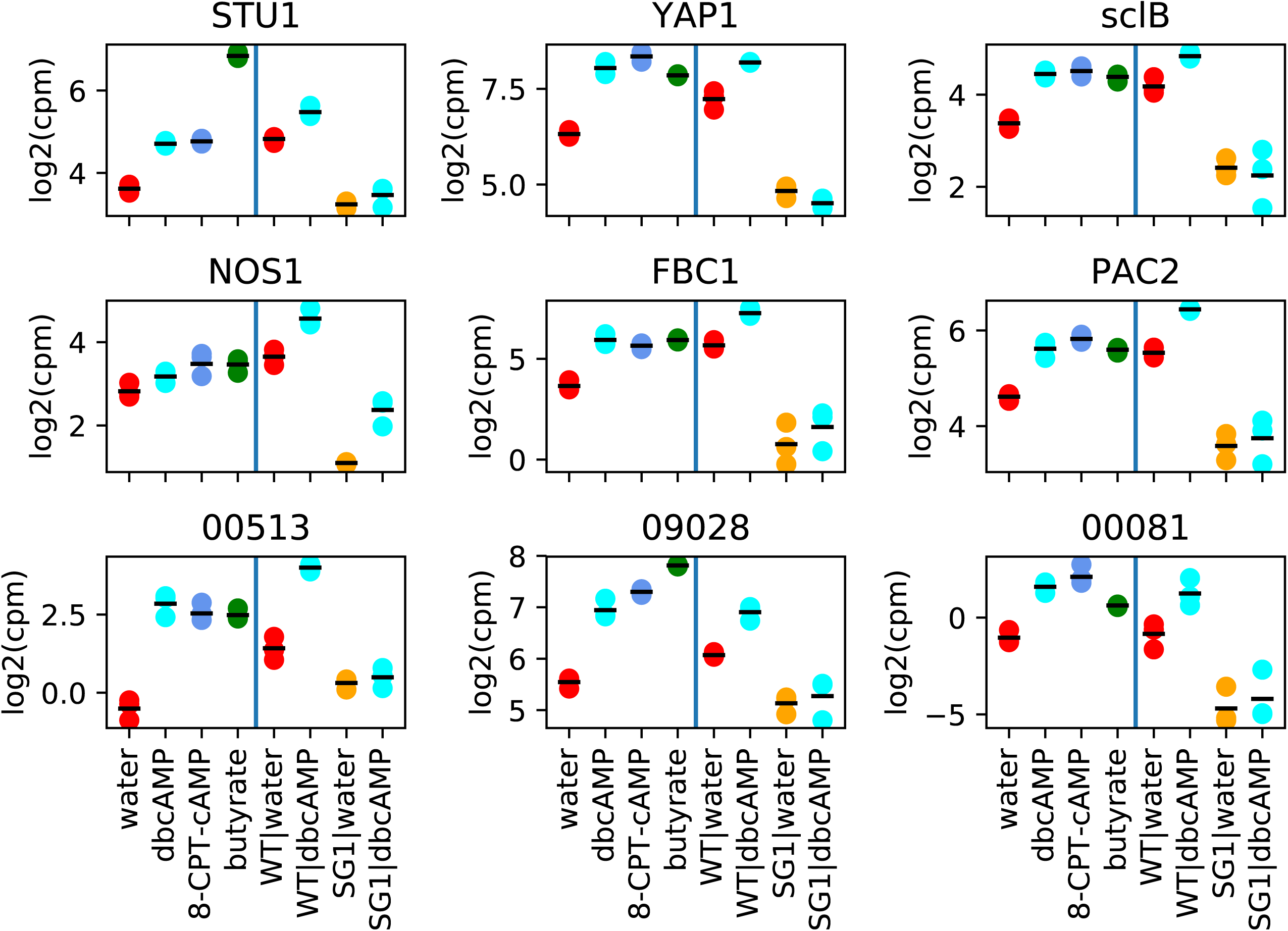
Transcript abundances of transcription factors in the stringently filamentous gene group (class 1 in Figure 3A), reported as log_2_ of read counts per million (cpm) in two separate experiments (as indicated by the two halves of each box). The conditions for the left half of each box are water, dbcAMP, 8-CPT-cAMP and butyrate for the wild-type strain. Conditions for the right half of each box are either water or dbcAMP for either the wild-type or SG1 strain, as indicated. Short black lines represent LIMMA fit values of transcript abundances for each condition. Each box is labeled with the name of the corresponding TF or a ucsf_hc.01_1.G217B transcript number for unannotated TFs (S5 Data of (3)).

**Table S1:** Strains used in this study

| Name | Referred in this study as | Comment |
| --- | --- | --- |
| **G217B-DA** | G217B | Derived from ATCC 26032 through passaging. |
| **G217Bura5-DA** | G217Bura5 or *ura5* | Derived from WU15 through passaging; Parental strain used to generate CRISPR mutants. |
| **G217Bura5 *msb2*::T-DNA** | SG1 | WU15 with T-DNA insertion upstream of *MSB2* ORF (11). |
| **G217Bura5-DA *fbc1*Δ** | *fbc1* | Deletion strain of *FBC1* (ATG+273..1555 deleted). |
| **G217Bura5-DA *pac2-1*** | *pac2* | Disruption strain of *PAC2* (1 bp insertion in position ATG+33). |
| **G217Bura5-DA *stu1-2*** | *stu1* | Disruption strain of *STU1* (2 bp insertion in position ATG+26). |
| **G217Bura5-DA *pACT1-tCATB*;**<br>***URA5*** | *ura5* + Overexpression control | Overexpression control. |
| **G217Bura5-DA *pACT1-FBC1-tCATB*;**<br>***URA5*** | *ura5* + *pACT1-FBC1* | Overexpression of *FBC1*. |
| **G217Bura5-DA pACT1-PAC2-tCATB; URA5** | *ura5 + pACT1-PAC2* | Overexpression of *PAC2*. |
| **G217Bura5-DA pACT1-STU1-tCATB; URA5** | *ura5 + pACT1-STU1* | Overexpression of *STU1*. |
| **G217Bura5-DA *fbc1*Δ pACT1-tCATB; URA5** | *fbc1 + Overexpression control* | Overexpression control in the *fbc1* strain. |
| **G217Bura5-DA *fbc1*Δ pACT1-FBC1-tCATB; URA5** | *fbc1 + pACT1-FBC1* | Overexpression of *FBC1* in the *fbc1* strain. |
| **G217Bura5-DA *fbc1*Δ pACT1-PAC2-tCATB; URA5** | *fbc1 + pACT1-PAC2* | Overexpression of *PAC2* in the *fbc1* strain. |
| **G217Bura5-DA *fbc1*Δ pACT1-STU1-tCATB; URA5** | *fbc1 + pACT1-STU1* | Overexpression of *STU1* in the *fbc1* strain. |
| **G217Bura5-DA *pac2-1* pACT1-tCATB; URA5** | *pac2 + Overexpression control* | Overexpression control in the *pac2* strain. |
| **G217Bura5-DA *pac2-1* pACT1-FBC1-tCATB; URA5** | *pac2 + pACT1-PAC2* | Overexpression of *FBC1* in the *pac2* strain. |
| **G217Bura5-DA *pac2-1* pACT1-PAC2-tCATB; URA5** | *pac2 + pACT1-PAC2* | Overexpression of *PAC2* in the *pac2* strain. |
| **G217Bura5-DA *pac2-1* pACT1-STU1-tCATB; URA5** | *pac2 + pACT1-STU1* | Overexpression of *STU1* in the *pac2* strain. |
| **G217Bura5-DA *stu1-2* pACT1-tCATB; URA5** | *stu1* + Overexpression control | Overexpression control in the *stu1* strain. |
| **G217Bura5-DA *stu1-2* pACT1-FBC1-tCATB; URA5** | *stu1* + pACT1-FBC1 | Overexpression of *FBC1* in the *stu1* strain. |
| **G217Bura5-DA *stu1-2* pACT1-PAC2-tCATB; URA5** | *stu1* + pACT1-PAC2 | Overexpression of *PAC2* in the *stu1* strain. |
| **G217Bura5-DA *stu1-2* pACT1-STU1-tCATB; URA5** | *stu1* + pACT1-STU1 | Overexpression of *STU1* in the *stu1* strain. |

**Table S2:**
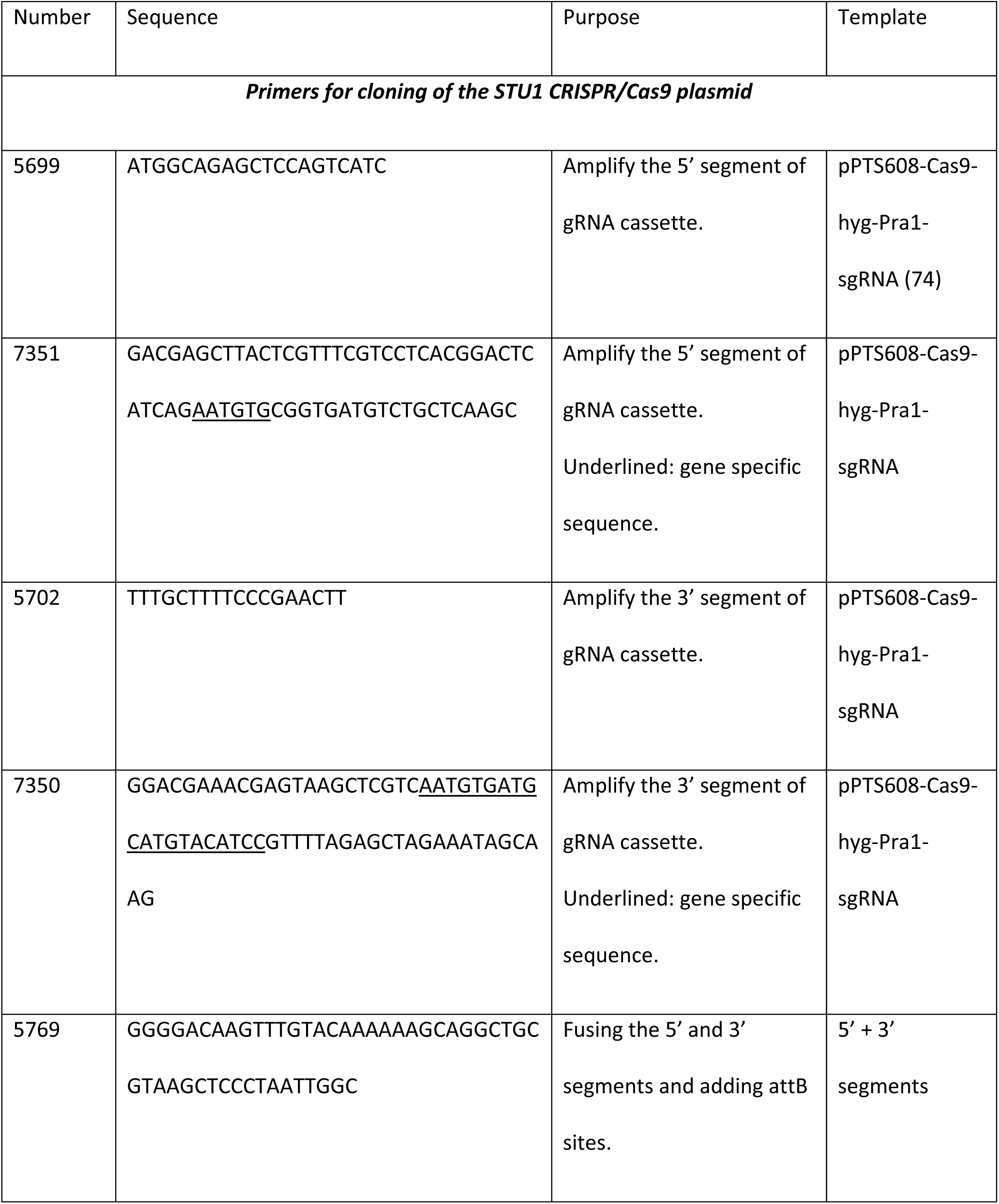
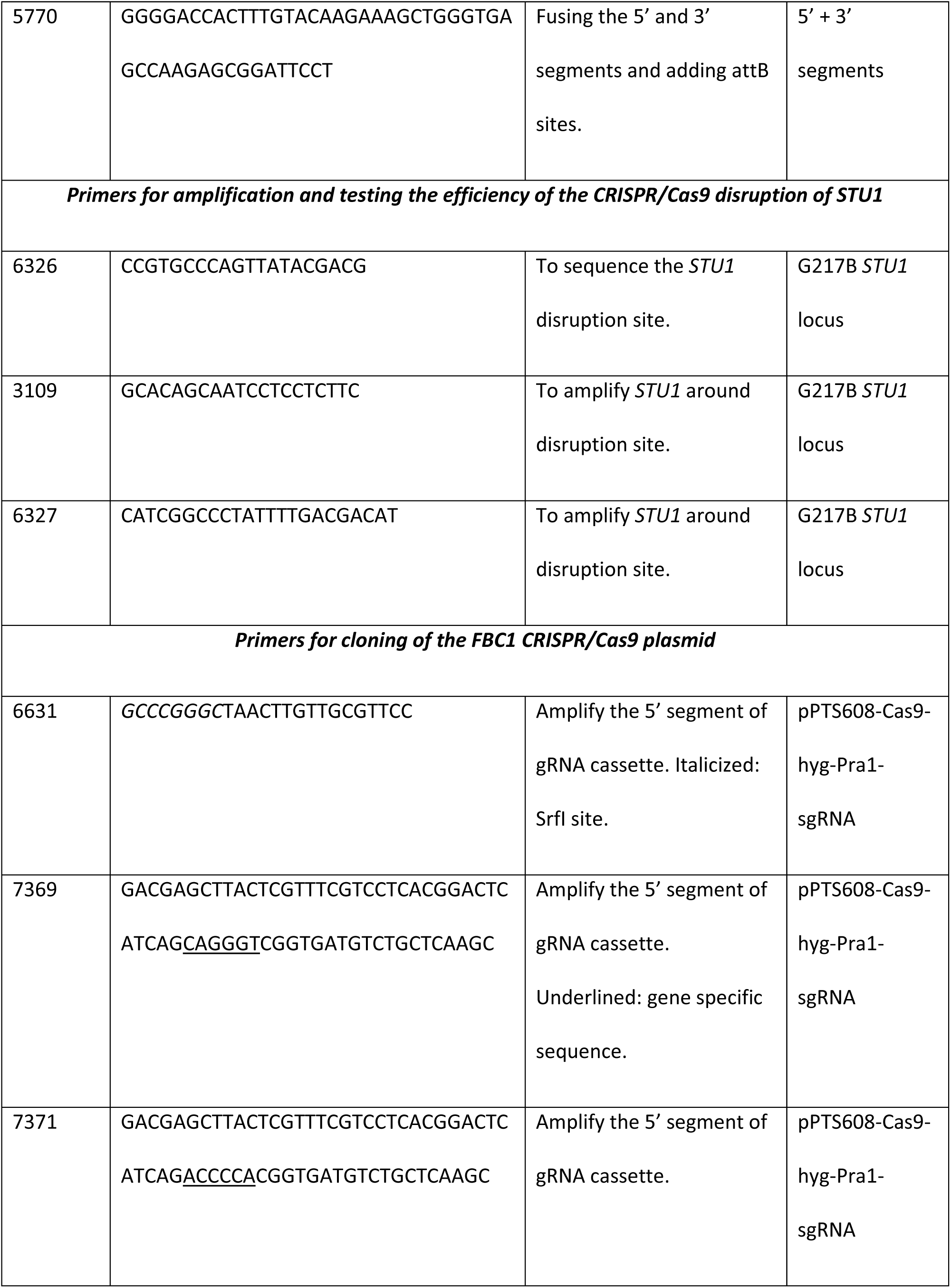
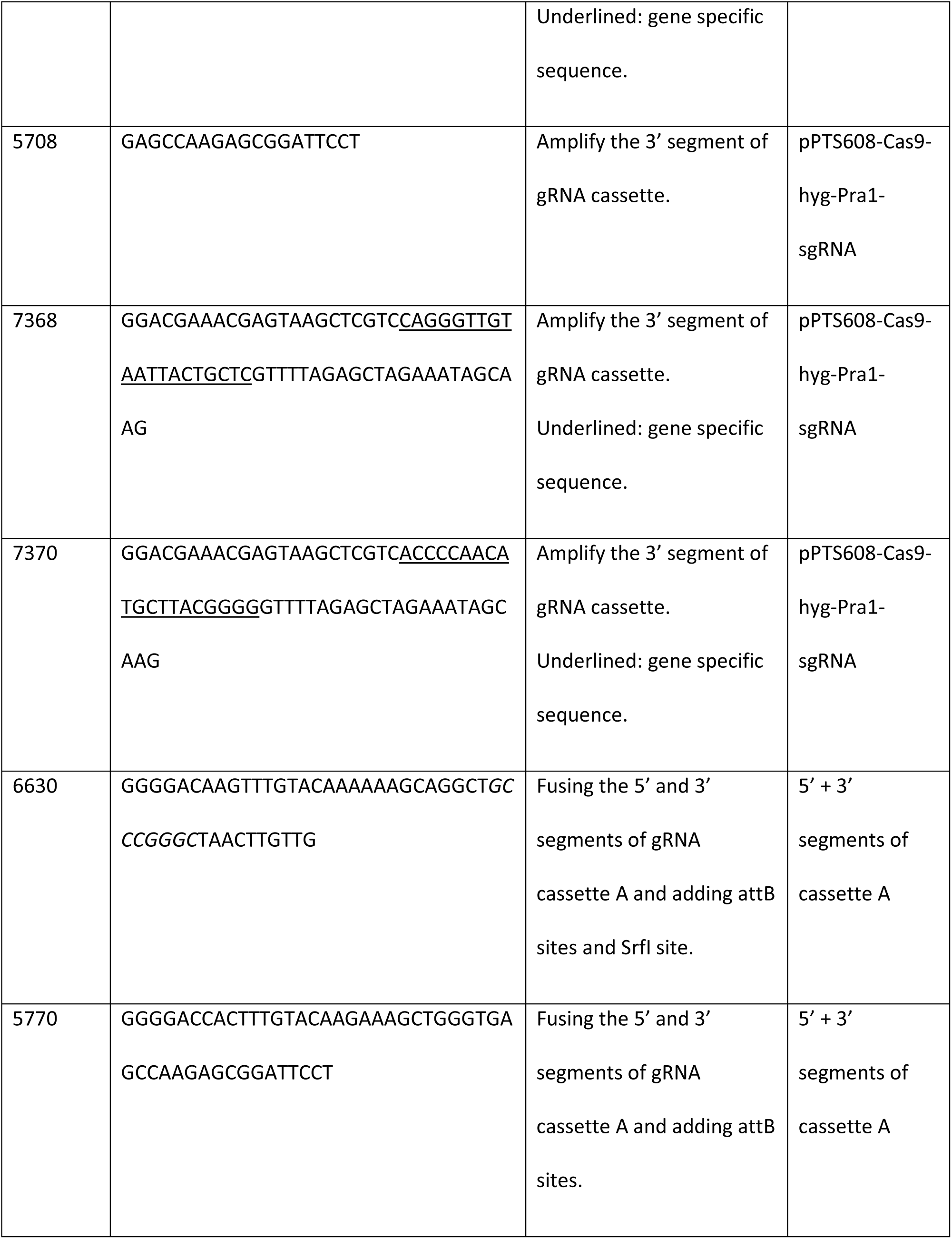
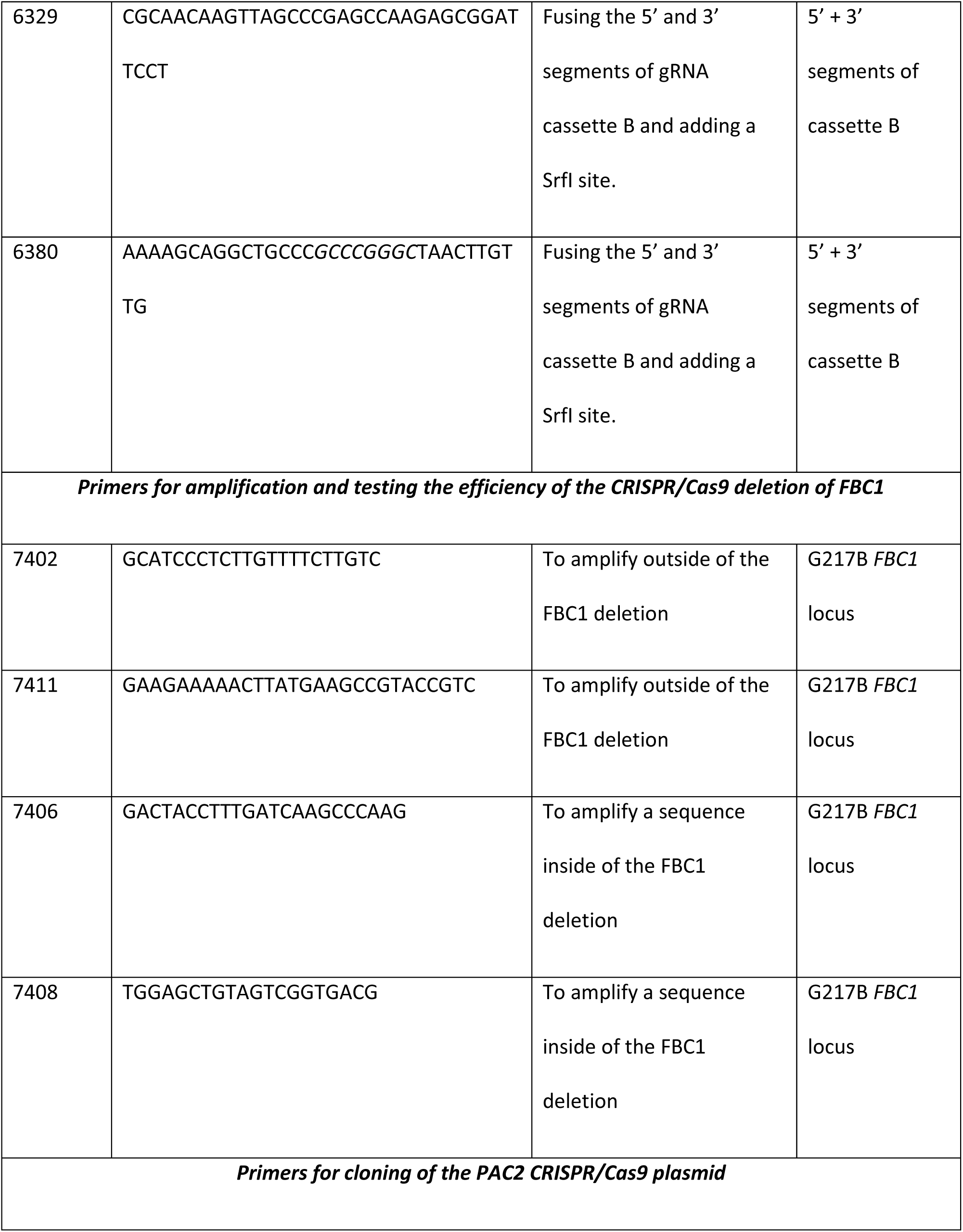
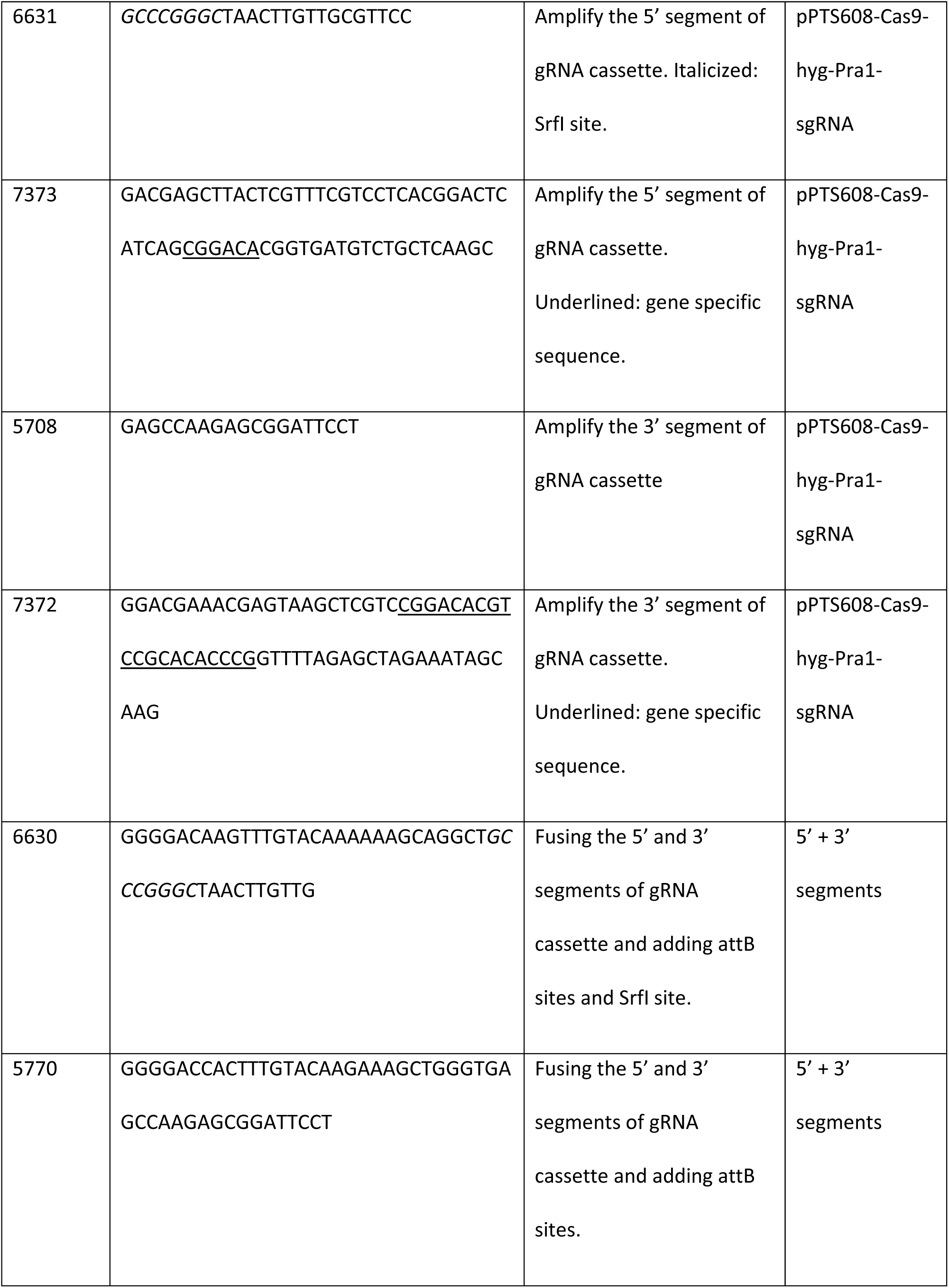
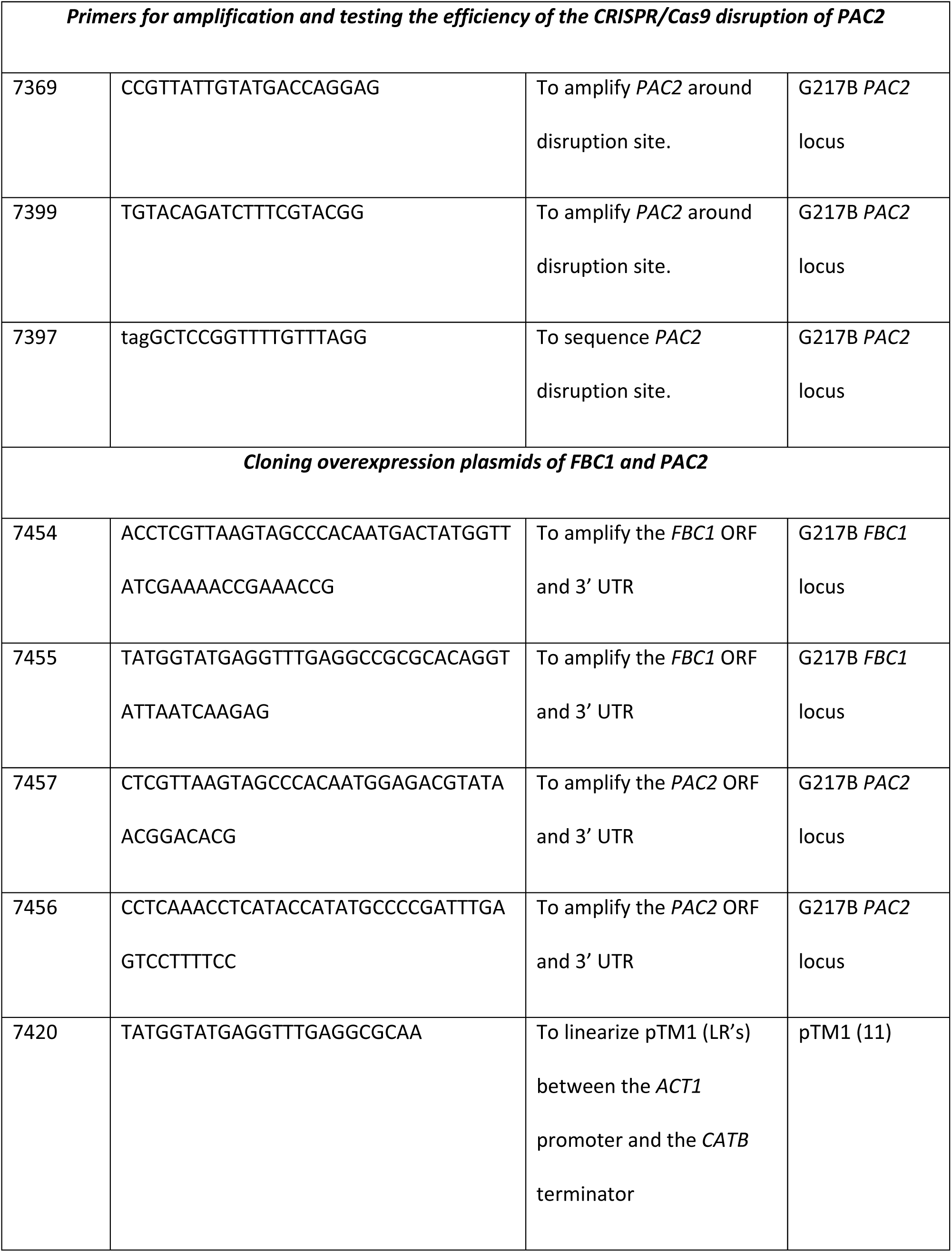
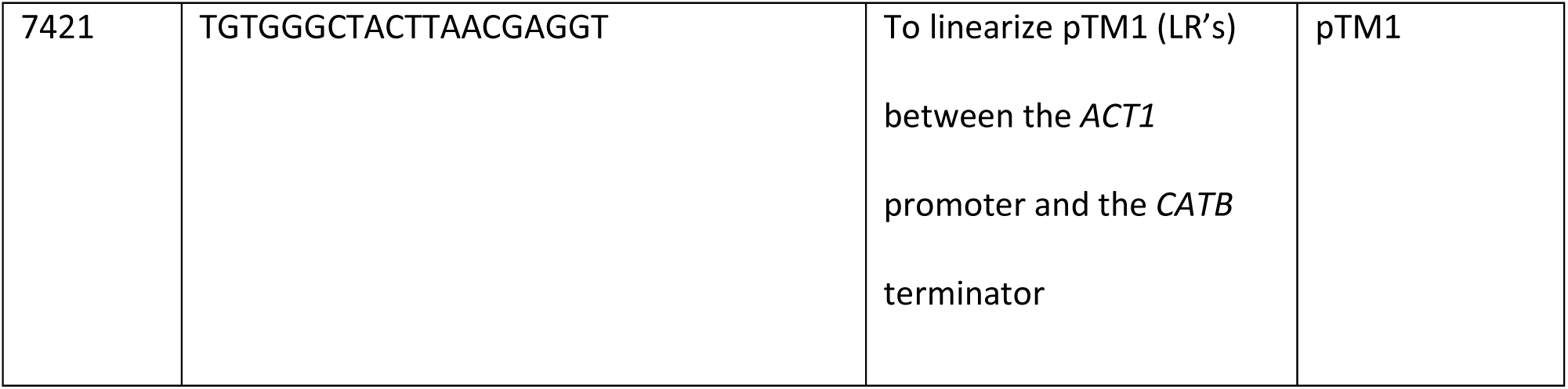
Primers used in this study

